# Repulsive interaction and secondary structure of highly charged proteins in regulating biomolecular condensation

**DOI:** 10.1101/2022.11.16.516834

**Authors:** Cheng Tan, Ai Niitsu, Yuji Sugita

## Abstract

Biomolecular condensation is involved in various cellular processes both functional and dysfunctional. Regulation of the condensation is thus crucial to avoid pathological protein aggregation and to maintain stable cellular environments. Recently, a class of highly charged intrinsically disordered proteins (IDPs), which are called the heat-resistant obscure (Hero) proteins, are shown to protect other client proteins from pathological aggregation. Besides the potential importance of this function, molecular mechanisms for how Hero proteins can protect other proteins from aggregation are not still known. Here we perform multiscale molecular dynamics (MD) simulations of Hero11, one of the Hero proteins, and the C-terminal region of TDP-43, as a target protein of Hero11, at various conditions to examine how they interact with each other. Based on the simulation results, three possible mechanisms have been proposed: (i) TDP-43 and Hero11 in dense phase reduces contacts with each other and shows faster diffusion due to the repulsive Hero11-Hero11 interactions, (ii) the amount of TDP-43 in dilute phase increases and their sizes become greater upon the attractive Hero11-TDP-43 interactions, and (iii) Hero-11 on the surface of small TDP-43 condensates avoids their fusions with the repulsive interactions. We also examine possible Hero-11 structures in atomistic and coarse-grained MD simulations and found disordered Hero-11 tend to assemble on the surface of the condensates, avoiding the droplet fusion effectively. The proposed mechanisms give us new insight into the regulation of biomolecular condensation in the cells and other conditions.

## Introduction

Proteins and nucleic acids can form biomolecular condensates via liquid-liquid phase separation (LLPS) [1]. Recently, LLPS has been observed in many cellular processes and considered as a general mechanism for the compartmentalization of biomolecules [2]. Formation of biomolecular condensations can be either functional or dysfunctional. The former includes genome organization [3,4], assembly of transcription machinery at super-enhancer [5], cellular signaling [6], and response to environmental stress [7], and the latter involves the assembly of pathogenetic ribonucleoprotein granules that leads to irreversible liquid-to-solid transitions [8]. The transactive response DNA binding protein 43 (TDP-43) [9] is one of the well-known examples that can form dysfunctional LLPS, whose consequent aggregation are related to several neurodegenerative diseases [9–11], including Limbic-predominant Age-related TDP-43 Encephalopathy (LATE) [12], Amyotrophic Lateral Sclerosis (ALS) [13], and Frontotemporal Dementia (FTD) [14]. Thus, understanding molecular mechanisms for how biomolecular condensations are regulated in the cell is crucial both in basic cellular biology and medical sciences.

Experimental studies have revealed many general molecular features in biological LLPS. Multivalency [15], usually driven by the proteins’ low complexity (LC) intrinsically disordered regions (IDRs) is one of the central principles underlying LLPS [16,17]. The C-terminal prion-like domain (PLD) of TDP-43 is a typical IDR that plays an essential role in TDP-43’s phase transitions [18,19]. Changes in the environmental conditions [20], mutations [21,22], or post-translational modifications (PTMs) such as acetylation [23,24] and phosphorylation [25] can regulate biological LLPS. Liquid droplets formed by RNA-binding proteins (RBPs) and RNAs can be modulated by RNA-protein ratio and length of RNA [26,27]. Interestingly, short bait RNAs composed of TDP-43 target sequences were used to prevent the neurotoxic aggregation of TDP-43 [19]. Similarly, a recent experimental study has identified a class of highly-charged intrinsically disordered proteins (IDPs), the heat-resistant obscure (Hero) proteins, that can protect proteins from pathological aggregation [28]. It was reported that TDP-43 aggregations can be suppressed by the co-expressed Hero proteins in cells [28]. Interestingly, shuffling of amino-acid sequence in Hero proteins does not change the activity of Hero proteins if the high fraction of charged residues is maintained [28]. The experimental results suggest the importance of charged residues in the anti-aggregation functions of Hero proteins, whereas their detailed molecular mechanisms are still unknown.

Theoretical and computational studies have been carried out to examine the phase behavior of IDPs for understanding the formation of LLPS at atomic details [29,30]. The stickers-and-spacers model provided a framework to describe the composition and interactions between multi-valent molecules [29,31,32]. The random phase approximation (RPA) theory was used to build connections between charge patterning and phase transition of IDPs [33,34]. However, most studies of the charged IDPs or polyampholytes focus on the promotion of phase separation due to the attractive interactions between proteins or between proteins and RNAs [33–36]. Improvements of all-atom force fields [37–39] and residue-level coarse-grained (CG) models [40–43] have been proven to be effective tools to investigate biomolecular interactions in dilute and dense phases. Notably, the hydrophobicity scale (HPS) CG model [40] has shown a great potential in capturing the sequence-specific thermodynamic properties [44–48]. Multiscale MD simulations have been applied to study the effects of a transient α-helical secondary structure [22,49] in TDP-43’s C-terminal IDR. They showed that the α-helical structure facilitates the formation of the dense phase and induces a higher critical temperature of TDP-43 [49].

In this study, we similarly perform multi-scale MD simulations of biomolecular condensates, while targeting on the anti-aggregation functions of Hero proteins. We select the C-terminal IDR of TDP-43 as proteins that form high-density condensates and examine how Hero11 can regulate the physical properties of condensations. The electrostatic repulsion between Hero proteins could be a key factor so that we also carried out simulations of a mixture of TDP-43’s C-terminal IDR and a chargeless mutant of Hero11. The secondary structure dependence of anti-aggregation function is also examined using atomistic and coarse-grained MD simulations. The repulsive interactions between Hero11s and the attractive interactions between TDP-43 and Hero11 regulate the formation of TDP-43 droplets and the kinetics of the protein. We propose three possible molecular mechanisms underlying the anti-aggregation function by Hero11 and discuss their general applicability to other Hero proteins as well as highly charged biomolecules like RNAs.

## Results

### Single chain simulations of TDP-43 and Hero11

To understand the phase behavior of TDP-43 and the anti-aggregation regulation function of Hero11, we performed atomistic and CG simulations of the C-terminal fraction of TDP-43 (residue 261-414, hereafter referred to as TDP-43 for convenience) and the full-length Hero11 of wild type (WT) and mutants (sequences listed in Supplementary Table S1). Previous studies have identified a short α-helical piece in TDP-43 and have shown that this secondary structure facilitates the phase separation of TDP-43 [22,49]. Interestingly, AlphaFold2 [50] predicts a structure of TDP-43 containing an α-helix in the same region. For Hero11, however, there are discrepancies in its secondary structure predictions between AlphaFold2 and the IDP-predicting tools, IUPred3 [51], flDPnn [52], and PONDR [53]. AlphaFold2 suggests two α-helices located in regions 17-25 and 38-72 of Hero11, while IUPred 3, flDPnn, and PONDR anticipate the whole chain with IDR propensities >0.5 (see Supplementary Figure S1).

To examine the secondary structure of Hero11, we employed the CHARMM36m force field [38] and carried out atomistic simulations of full-length Hero11 in explicit solvent (see Methods for detailed simulation conditions). We started from two different initial structures. One was downloaded from the AlphaFold Protein Structure Database [50] (entry Q9UNZ5) and contains two α-helices. The other is a random coil reconstructed from a short CG simulation using the HPS model (Supplementary Figures S2A and S2B). After energy minimization and equilibration, we performed ∼1μs simulations for each system. In the simulation starting from AlphaFold2-predicted structure, the first α-helix (residue 17-25) unfolds after ∼0.6μs, while the second α-helix is only partly sustained (residue 38-55, see Supplementary Figure S2). In the simulations starting from a random coil, a short piece of α-helical structure at residue 44-50 emerges in the early stage and is preserved until the end. Structures sampled from these two simulations are pretty different, showing that 1μs simulation is insufficient to achieve the equilibrium. To obtain statistically more reliable results, it is necessary to perform much longer MD simulations (> 100 μs) or use one of the enhanced conformational sampling algorithms for exploring protein conformational landscapes [54–56]. Regardless of initial structures, a short region (residue 44-50) is predicted as a helical structure in all the MD simulations, suggesting that Hero-11 cannot be a completely random structure in solution.

We next utilized CG simulations to explore the conformations of Hero11 and TDP-43. Since our atomistic simulations with a classic force field did not provide a substantial conclusion about the secondary structure of Hero11, we decided to model two extreme cases: one harboring the two α-helices predicted by AlphaFold2 and the other with the entire peptide as an IDP. Here, we employed the HPS potentials [40] and the HPS-Urry parameter set [44] to define the interactions in the IDR. For the α-helical regions, we used the AICG2+ model [57] to restrain the folded secondary structures. Figure 1 shows the sequences and secondary structures of Hero11 and TDP-43.

**Figure 1.**
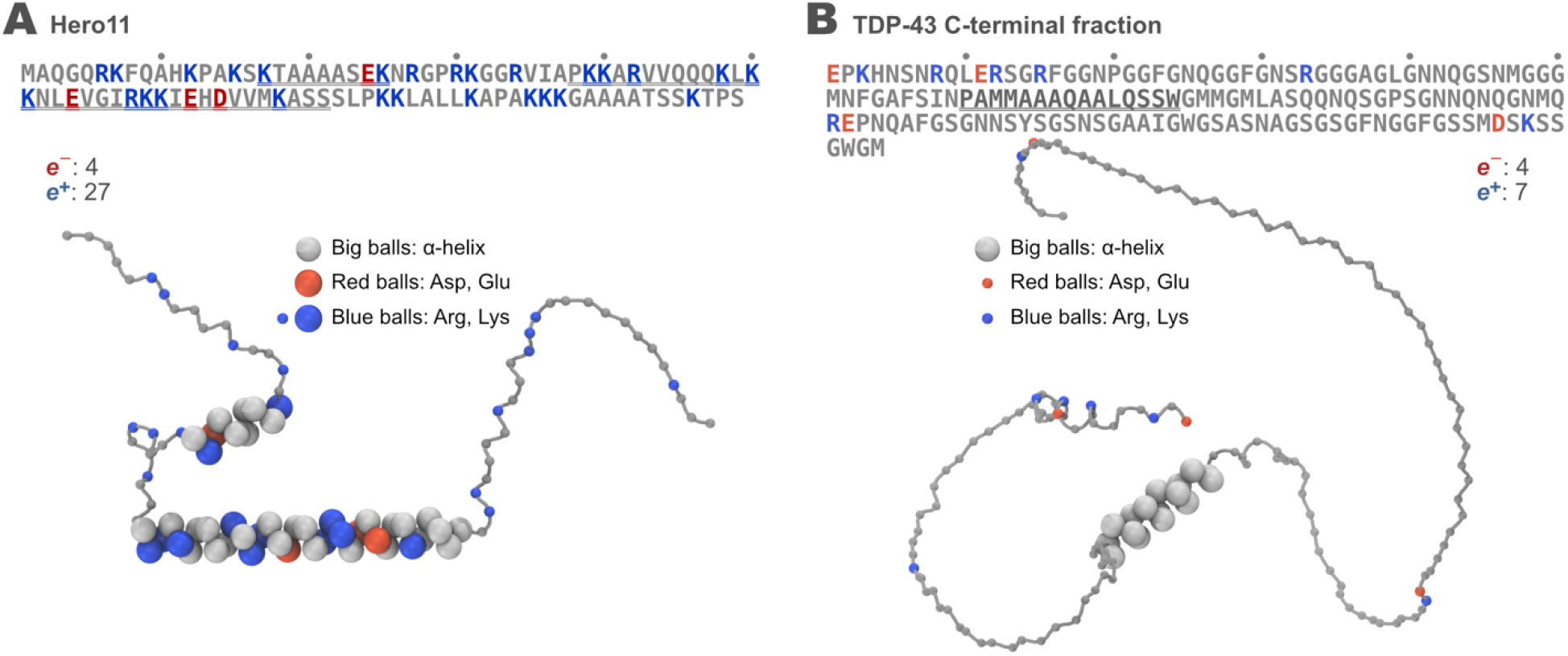
Sequence and coarse-grained modeling of Hero11 and TDP-43 C-terminal fraction. Each amino acid residue is represented by one bead in the simulations. Particles are shown in different sizes to represent different treatments: small particles are modeled with the HPS, whereas large particles are modeled with the AICG2+ to maintain the α-helical structures based on AlphaFold2 predictions.

We first conducted simulations of a single-chain TDP-43 at temperatures ranging from 250K to 350K. At all temperatures, the α-helices are well maintained during the simulations (see Supplementary Figure S3B). The IDR tails of TDP-43 are flexible, and the whole structure shows an increasing in radius-of-gyration (R_*g*_) as temperature increases (Supplementary Figure S3C). These results show that our method of combining AICG2+ and HPS models can be used to study the conformations of proteins consisting of both folded domains and IDRs. Using the same method, we also simulated a single-chain Hero11-WT with the α-helical structure (named Hero11-WT-α) or without any α-helical structure (named Hero11-WT-noα), respectively. Contact analysis demonstrates that when applied, the AICG2+ potentials preserve the folded structures of α-helices (Supplementary Figure S4B). Interestingly, we find that Hero11-WT-α has larger R_*g*_ values than Hero11-WT-noα at all the simulated temperatures (Supplementary Figure S4C). This result shows that the α-helical secondary structures result in relatively more extended conformations of Hero11-WT.

### Phase behavior of wild-type Hero11

Before applying our model to multiple-chain simulations of Hero11, we further validated our method on TDP-43. Following the slab simulation strategy [40], we generated a system composed of 100 chains of TDP-43 in a 180Å × 180Å × 3000Å box with the periodic boundary condition. The system was simulated at ten different temperatures from 260K to 350K with a 10K interval. During the simulations, we monitored the protein density distribution along the *z*-axis (the longest dimension). Practically, we divided the z-axis into small bins of size Δ_*z*_ = 30Å and computed the local density of TDP-43 molecules in each bin (more details are explained in Methods). Based on the density analysis, we then determined the dense and dilute phases at temperatures lower than the critical temperature (*T*_*C*_). In Figure 2A, we show a plot of the time evolution of TDP-43’s density (represented by the intensity of the blue color) in a simulation at 295K. A representative structure of the simulation box around the dense phase is shown in Figure 2B. In the dense phase, TDP-43 forms extensive intra- and inter-molecular interactions. In the dilute phase, TDP-43 chains are primarily free and seldom form contacts with each other. The density data at different temperatures are then used to plot the phase diagram of TDP-43, which is shown in Figure 2C (solid line and dots). As a control, we also built a model of TDP-43 without holding the α-helical structure and carried out the same simulation and analysis. Our results show that the α-helical secondary structures raise the critical temperature of TDP-43 by ∼5K (Figure 2C). Detailed contact analysis reveals that the α-helical region contributes more inter-chain interactions than the IDRs (Supplementary Figure S5). These results are consistent with the previous study [49], although the modeling methods and parameters differ.

**Figure 2.**
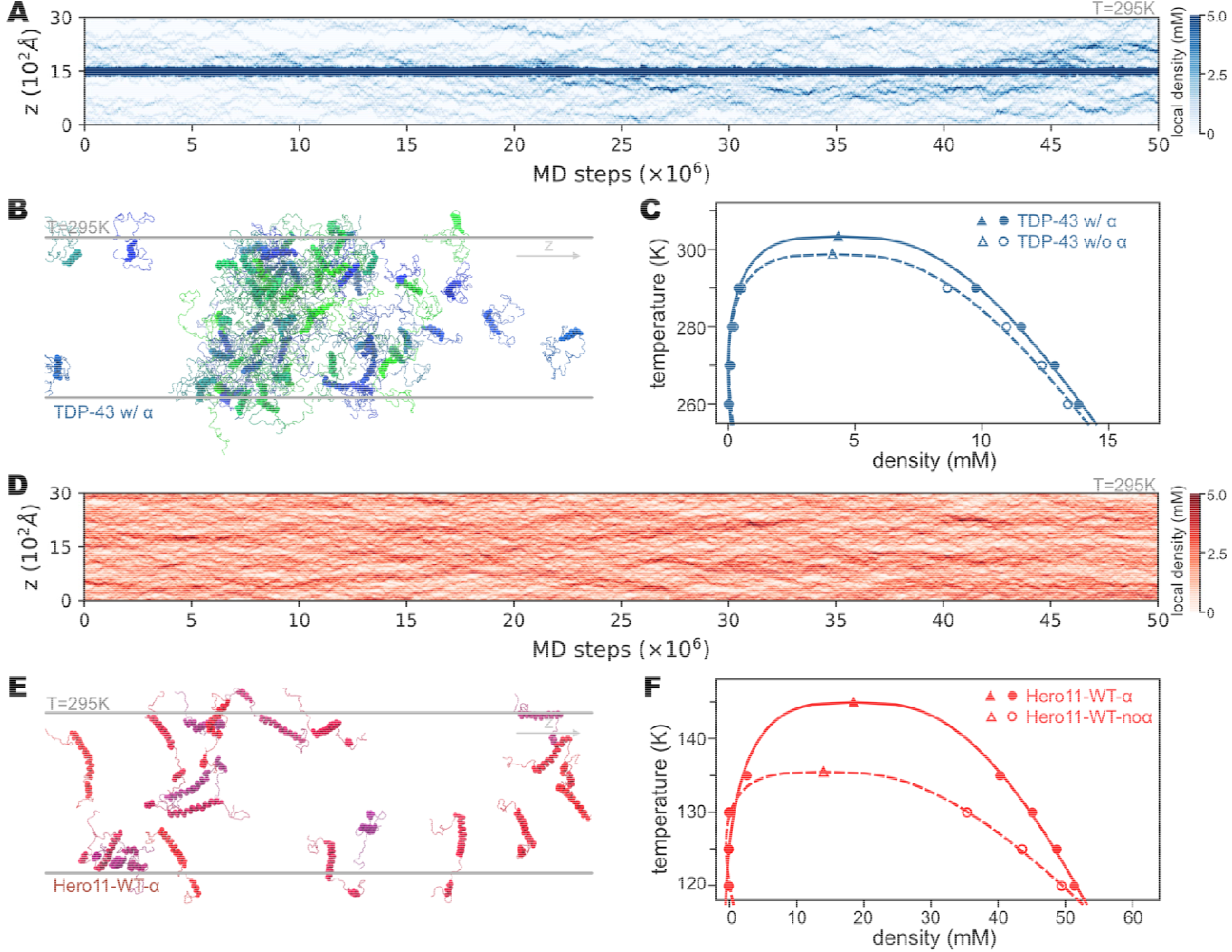
Multi-temperature slab simulations of homotypic TDP-43 and Hero11-WT. (A) Time evolution of TDP-43 local density along the z-axis (the longest dimension of the simulation box). In each simulation, 100 chains of TDP-43 are put in a box of. The z dimension is divided into 100 bins with length and local density is calculated in each bin. Intensity of the blue color represents local density, as indicated by the color bar. (B) The last structure of a -step slab simulation of TDP-43 at 300K. For clarity, we only show the region around the condensate of TDP-43. Chains are shown in different colors to distinguish them. (C) Phase diagram of TDP-43 with (solid line and dots) and without (dashed line and empty dots) the α-helical structure modeled by AICG2+. Circles represent the simulated densities and triangles represent the fitted values (see Methods for details). (D) is the same as (A) but for Hero11-WT. Density is represented by the intensity of red, as indicated by the color bar. (E) Part of the last structure of a -step slab simulation of Hero11-WT at 300K. Chains are shown in different colors. (F) is the same as (C) but for Hero11-WT-α and Hero11-WT-noα.

We next conducted CG simulations for multiple chains of Hero11-WT-α and Hero11-WT-noα, respectively. Hero11-WT has many positively charged residues (Figure 1A), and the net charge of a single chain is +23. Experiments have shown that such sequence feature results in a lower tendency to form condensates than typical LLPS or aggregation-prone proteins such as TDP-43 [28]. Consistently, our simulation results connect the sequence of Hero11-WT with its phase behavior. Figure 2D shows the density profile of Hero11-WT-α simulated at 295K. Hero11-WT-α does not assemble into a stable concentrated phase, as TDP-43 does at the same temperature (compare with Figure 2A). A snapshot of the Hero11-WT-α structure at 295K is shown in Figure 2E, demonstrating an unconsolidated structure with rare inter-chain contacts.

We then decreased the simulation temperature to determine the *T*_*C*_ of Hero11-WT. Being modeled as a pure IDP, Hero11-WT-noα shows a typical liquid-like condensation as temperature decreases from 140K to 120K (Supplementary Figure S6A-C). In contrast, with the α-helical structures maintained, Hero11-WT-α forms liquid-crystal-like smectic phases at temperatures ∼140K. When the temperature drops to 120K, the size of Hero11-WT-α cluster and the anisotropic orientational ordering increase (Supplementary Figure S6D-F). Since our model is not designed to study the liquid-crystal-like phase behaviors, we decide not to concentrate too much on this phenomenon in the current work. Nevertheless, we plot the phase diagrams of Hero11-WT-α and Hero11-WT-noα in Figure 2F. Interestingly, the tendency that α-helical structures yield a relatively higher *T*_*C*_ than the pure IDR model [49] also holds for Hero11-WT. At low temperatures, Hero11-WT-α also uses its α-helical region to form extensive inter-chain interactions (Supplementary Figure S7). Interestingly, regions with a higher charge density (for example, the K_85_K_86_K_87_ piece) have lower probabilities of forming inter-chain contacts in both Hero11-WT-α and Hero11-WT-noα (Supplementary Figure S7). As discussed earlier, the actual behavior of Hero11-WT can be viewed as an intermediate between Hero11-WT-α and Hero11-WT-noα. Therefore, our simulations indicate that Hero11-WT bears a much lower *T*_*C*_ than LLPS-prone IDPs like TDP-43. These results provide molecule-level structural evidence for previously discovered solubility of the Hero proteins [28].

Previous simulation studies have investigated the *R*_*g*_ of IDR in the condensate and its relationship with conformational entropy gain during phase transition [58,59]. Inspired by these works, we also calculated the *R*_*g*_ of TDP-43 and Hero11-WT from our simulations (Supplementary Figure S8). Consistent with earlier findings [58,59], all the proteins simulated here have larger *R*_*g*_ values in the condensate than in the dilute phase. While the single-chain *R*_*g*_ grows monotonically as temperature, the dense phase *R*_*g*_ is almost independent of temperature. Interestingly, Hero11-WT-α has larger *R*_*g*_ values than Hero11-WT-noα in the condensates, although the difference between them is smaller than the single-chain results (Supplementary Figure S8B).

### Phase behavior of Hero11 mutants

Mutation experiments has suggested that the positive charges on Hero11-WT dominates its phase behavior [28]. Following the experimental design, we also simulated several Hero11 mutants, including the “KRless” mutant, where all the Lysine and Arginine residues are mutated to Glycine, and the “scrambles”, in which Hero11’s sequence is randomly shuffled [28]. A complete list of all the mutant sequences can be found in Supplementary Table S1.

Similar to the case of Hero11-WT, we first ran all-atom simulations for Hero11-KRless using the CHARMM36m force field [38]. Two initial structures are used, either from the AlphaFold2 predicted structure or a random coil. During the 1μs simulation of the predicted structure, we find that the α-helix is partially unfolded, and only a short part (residue 43-55) is kept (Supplementary Figure S2C). In the simulation from a random coil, some short α-helix and β-strand are formed transiently but are not stable (Supplementary Figure S2D). Similar to Hero11-WT, the Hero11-KRless simulations also did not present convergent results. Therefore, we consider the two extreme conformations in the CG simulations, Hero11-KRless-α and Hero11-KRless-noα, respectively.

We performed CG simulations for systems containing 100 chains of Hero11-KRless-α or Hero11-KRless-noα at temperatures from 210K to 300K. Using the same analysis method as for Hero11-WT, we plot the time evolution of density along the *z*-axis, as shown in Figure 3A. Unlike Hero11-WT, Hero11-KRless-α forms a stable cluster at 250K, with dynamic dissociation and association of single chains. Figure 3B shows a representative snapshot of the simulation box of Hero11-KRless-α at 250K. As can be seen, α-helical structures still contribute a large number of interactions in the dense phase of Hero11-KRless-α, as indicated by the contact analysis (Supplementary Figure S9). We then use the density and temperature data to plot the phase diagram for the KRless mutants, as shown in Figure 3C (gray dots and lines). Interestingly, we also observe an elevated *T*_*C*_ for the model with the α-helical region (Hero11-KRless-α) than the one with no α-helix (Hero11-KRless-noα), suggesting the effect of α-helix in biomolecular condensate to be a general mechanism. Rather importantly, it is clear that removing the positive charges from the sequence significantly raises the Hero11’s critical temperature (Figure 3C). In other words, the electrostatic repulsion due to the high content of positively charged residues in Hero11-WT dominates its phase behavior. To further testify to this conclusion, we mutate all the charged residues (including 27 positive and 4 negative residues, see Figure 1A) in Hero11 and model them as Hero11-chargeless-α and Hero11-chargeless-noα, respectively. By carrying out simulations at multiple temperatures, we get the phase diagrams for these two chargeless mutants (see Supplementary Figure S10). Compared with the Hero11-KRless mutants, which have a net charge of −4*e*, the Hero11-chargeless mutants are neutralized and show even higher *T*_*C*_ than the Hero11-KRless mutants. These results confirm the hypothesis that inter-chain electrostatic interactions between the large amount of positively charged residues are the main reason that Hero11 does not form condensate at physiological temperatures [28].

**Figure 3.**
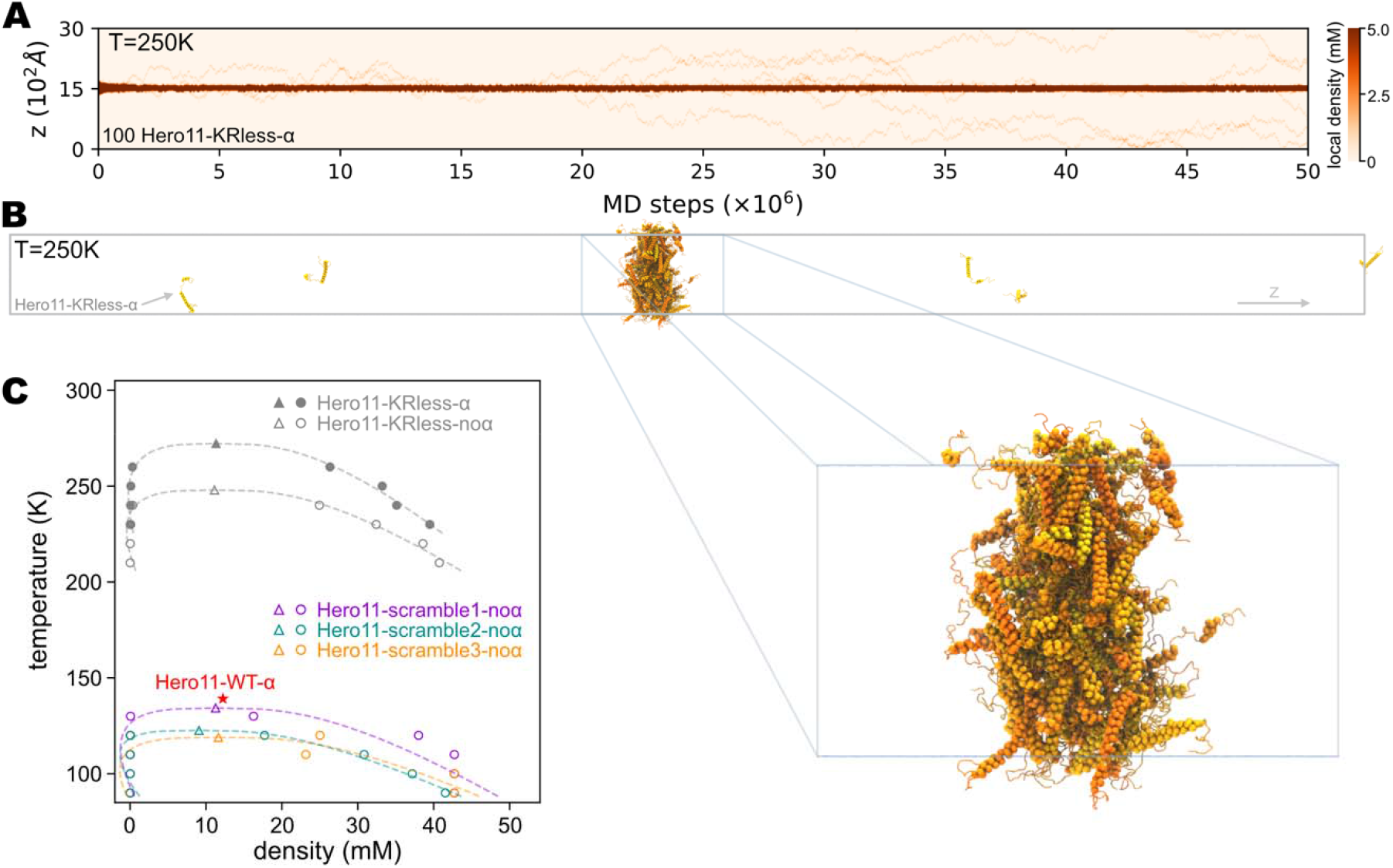
Phase behavior of Hero11 mutants. (A) A representative graph of local density of Hero11-KRless mutant over time. Intensity of the orange color represents local density, as indicated by the color bar. Local density has the same definition as Figure 2A. (B) The last structure of a 5 × 10^7^-step simulation of Hero11-KRless at 250K. Colors from yellow to orange are used to distinguish different chains. (C) Phase diagram of Hero11-scrambles (see Supplementary Table S1 for sequences) and Hero11-KRless mutants. Circles are simulated values and triangles are fitted values (see Methods for details). The red star represents the wild type (WT) Hero11’s critical point.

In addition to the KRless and chargeless mutants, in which the fraction of charges is altered, we also simulated the mutations where the charge fraction is kept, but the sequence is randomly shuffled (named as Hero11-scramble*N, N* = 1, 2, 3). We modeled these mutants as pure IDPs since there is no reason to assume the α-helices to be preserved for the randomly shuffled sequences. We plot the phase diagrams for these mutants in Figure 3C. As can be seen, all the three tested scrambles show similar *T*_*C*_ values as Hero11-WT-noα. These results agree with previous experiments showing that the charge fraction is more crucial than the amino-acid sequence in determining the phase behavior of the Hero proteins [28].

### Regulation of TDP-43 condensate by Hero11

Next, we explored the mechanism for Hero11’s anti-aggregation effect on TDP-43, which was found in the experiments [28]. Here, using the slab method, we simulated a system consisting of 100 Hero11-WT-α and 100 TDP-43. As described in Methods, we prepared a consolidated mixture of Hero11-WT-α and TDP-43 as the initial structure. Simulations were performed at ten different temperatures ranging from 260K to 350K. Figures 4A and 4B show the time evolution of the densities of TDP-43 and Hero11-WT-α at two temperatures, 250K and 295K, respectively. As can be seen, density profiles of the two proteins are highly coupled. At 250K, TDP-43 and Hero11-WT-α chains jointly form the high-density condensate, as indicated by the darkest blue and red regions (Figure 4A). In contrast, at 295K, the dense phase formed by TDP-43 and Hero11-WT-α begins to dissolve soon after the beginning of simulation. The largest cluster diverges into several smaller ones in the last ∼2.5 × 10^7^ steps (Figure 4B). The last structures of these simulations are shown in Figure 4C (*T* = 250*K*) and 4D (*T* = 295*K*), respectively. It can be seen that at 250K, some Hero11-WT-α chains join the condensate formed by TDP-43. At 295K, Hero11-WT-α still forms contacts with TDP-43, but a clear boundary between the two phases disappears. Compared with the homotypic TDP-43 system at 295K (Figure 2A), these results illustrate how Hero11-WT-α changes the behavior of TDP-43. We then use the density data at 260K, 270K, 280K, and 290K to plot the phase diagram of TDP-43 in the presence of Hero11-WT-α. As shown in Figure 4E (cyan line), adding Hero11-WT-α decreases the *T*_*C*_ of TDP-43.

**Figure 4.**
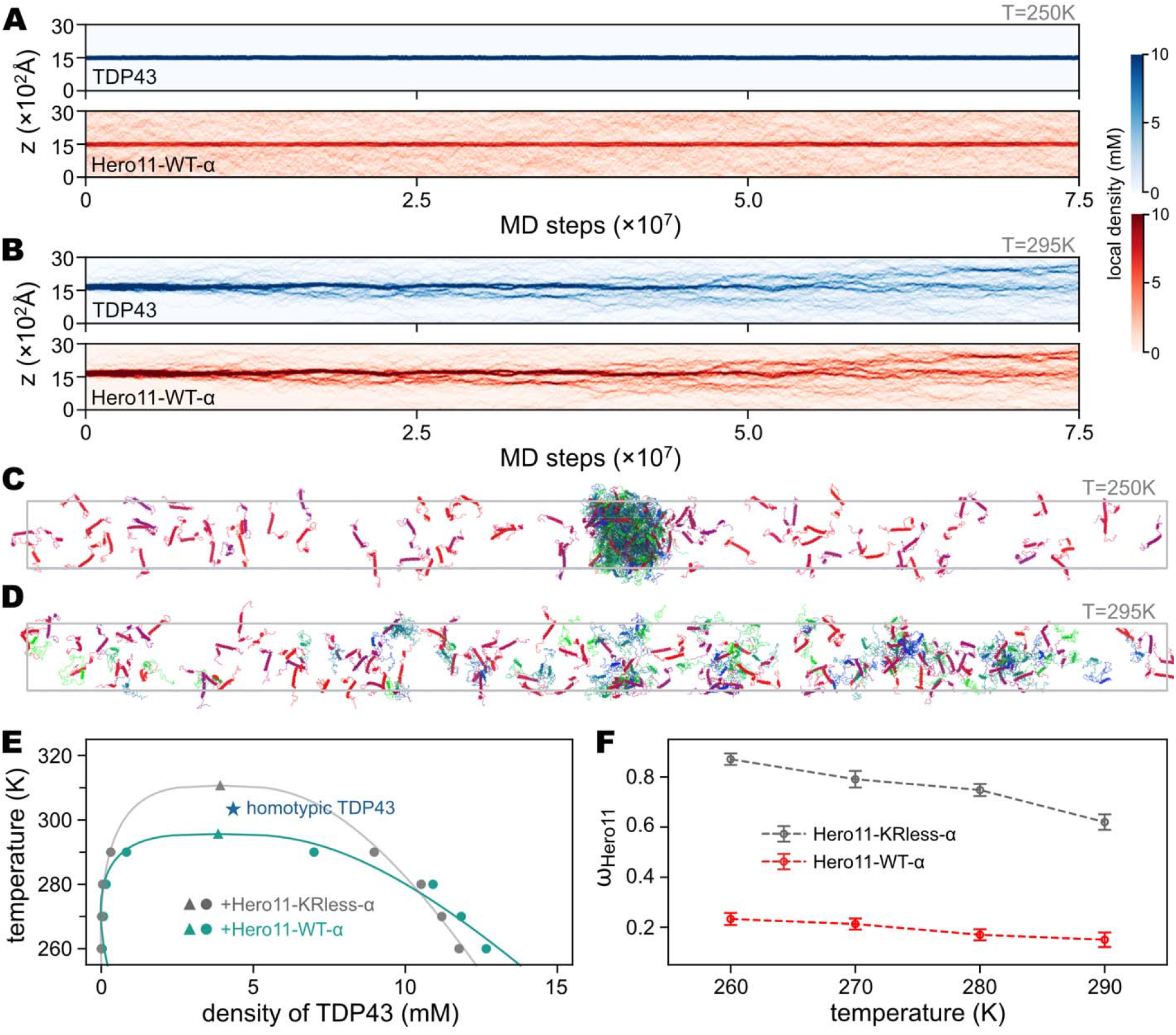
Slab simulations of heterotypic systems consisting of 100 TDP-43 and 100 Hero11. (A) and (B) show representative local density profiles of TDP-43 and Hero11-WT-α simulated at 250K and 300K, respectively. Intensity of blue (red) represents local density of TDP-43 (Hero11-WT-α), as indicated by the color bars. (C) and (D) show the last structures of the 250K and 300K trajectories, respectively. (E) Phase diagram of TDP-43 in the presence of Hero11-WT-α (cyan) or Hero11-KRless-α (gray). Circles represent the quantities calculated from the simulations, whereas triangles are the fitted values (see Methods). (F) Fraction of Hero11 located in the condensate of TDP-43 (*ω*_*Hero*11_) as a function of simulation temperature. For (E) and (F), only the last 1/3 of each simulation is used for analysis.

As shown in Figure 3, the mutations from Lys/Arg to Gly drastically change the phase behavior of the homotypic Hero11. We then wonder how these mutations affect the co-condensation of Hero11 and TDP-43. From simulations of the mixture of 100 Hero11-KRless-α and 100 TDP-43, we find that at 295K, the dense phase formed by both proteins is stably maintained (Supplementary Figure S11). In Figure 4E, we plot the phase diagram of TDP-43 from the simulations where Hero11-KRless-α is present. As can be seen, without the high content of positively charged residues, Hero11-KRless-α promotes *T*_*C*_ of TDP-43 to ∼310K (Figure 4E). Removing all the charged residues, Hero11-chargeless-α raises TDP-43’s *T*_*C*_ to a higher value of ∼320K (see Supplementary Figure S12A). Interestingly, we find that both Hero11-WT-α and Hero11-KRless-α enter the condensate of TDP-43, following a previously proposed “scaffold and client” model [60–62]. However, the fraction of “client”-Hero11 (*ω*_*Hero*11_) is smaller in the case of Hero11-WT-α than that of Hero11-KRless-α, due to the strong inter-chain repulsion (Figure 4F). These results show that the electrostatic interactions between positively charged residues in Hero11-WT-α determine its regulatory functions on the phase behavior of the other proteins.

### Mechanism underlying the Hero11’s anti-aggregation function

The simulation results of heterotypic systems consisting of TDP-43 and Hero11 are in good agreement with the previous experimental results [28]. Next, we go into more detailed mechanisms for Hero11’s anti-aggregation function by performing simulations of heterotypic systems composed of 100 TDP-43s and variant numbers (10*n, n* = 1, 2, …, 9) of Hero11-WT-αs. By analyzing densities of TDP-43 and Hero11-WT-α in different phases and at different temperatures, we plot the binary phase diagram in Figure 5A. We use data from systems including 20, 40, 60, 80, and 100 Hero11-WT-αs simulated at temperatures of 260K (red), 270K (green), 280K (blue), and 290K (purple), respectively. The dotted tie-lines connect the densities of the dilute phase (empty dots) and the dense phase (solid dots) from the same simulation. The tie-lines intersect at a single point for systems with the same number of Hero11-WT-α, corresponding to the averaged densities of proteins in the whole simulation box. As a reference, we also show the density of TDP-43 analyzed from the simulations of homotypic TDP-43 (“+” and “x” dots in Figure 5A). As can be seen, as the temperature increases, the biphasic region shrinks, suggesting an upper consolute temperature (UCT) above which no phase separation happens.

**Figure 5.**
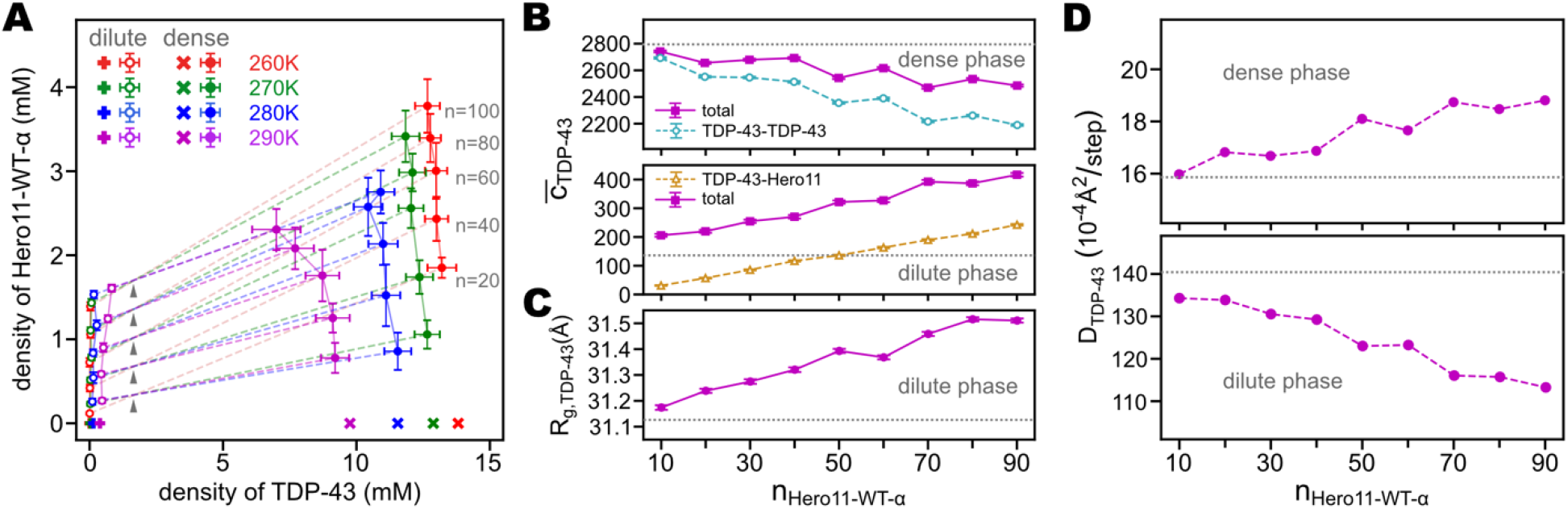
Effects of Hero11-WT-α on TDP-43 condensation. (A) Binary phase diagram of TDP-43 and Hero11-WT-α. Density coordinates calculated from the same simulations are connected by dashed lines. Note that simulations with the same number of Hero11-WT-α intersect at the same point (indicated by gray triangles). The number of Hero11-WT-α corresponding to each triangle are *n* = 20, 40, 60, 80, and 100, from lower to upper, as indicated by the numbers next to the high-density dots for 260K (red). Color of lines and dots represent the simulation temperatures, as indicated by the legends. “x” and “+” symbols represent the densities in the dense and dilute phase of the homotypic TDP-43 system, respectively. Error bars represent standard deviations. (B) Average number of contacts formed by any TDP-43 chain 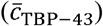 with other chains of TDP-43 or Hero11-WT-α. Upper and lower panels are for TDP-43s in the dense and dilute phases, respectively. The total number of 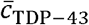 (purple) is the sum of TDP-43-TDP-43 (cyan) and TDP-43-Hero11 (orange) contacts. For clarity, in the dense phase we don’t show TDP-43-Hero11 data, while in the dilute phase TDP-43-TDP-43 is hidden. Horizontal gray lines represent the contact numbers calculated in the homotypic TDP-43 systems. (C) Radius of gyration of TDP-43 (*R*_*g*,TDP − 43_) in the dilute phase as a function of the number of Hero11-WT-α included in the simulation (*n*_Hero11-WT-α_). The horizontal dotted line represents *R*_*g*_ value of TDP-43 calculated from single-chain simulations. Error bars in (B) and (C) represent standard errors. (D) 3-dimensional diffusion coefficient of TDP-43 (*D*_TDP − 43_) in the dense (upper) and dilute (lower) phases as a function of *n*_Hero11-WT-α_. Horizontal dotted lines show the values of *D*_TDP − 43_ calculated from the simulations of homotypic TDP-43. Results in (B), (C), and (D) are based on simulations at temperature 290K.

We then focus on the density changes caused by adding Hero-WT-α to the TDP-43 condensate. Compared with the densities of the homotypic TDP-43 system, adding Hero11-WT-α loosens the dense phase, as indicated by the high-density branches in Figure 5A. This effect is more prominent at higher temperatures (290K) than at lower temperatures (280K to 260K). A larger number of Hero11-WT-α also loosens the dense phase more, as seen from the concave curve of the high-density branch at 290K in Figure 5A. We consider this phenomenon nontrivial since the “client” proteins do not necessarily decrease the density of condensates formed by the “scaffold” proteins [32,47,62]. On the other hand, our results also show that Hero11-WT-α increases the saturation concentration of TDP-43, as indicated by the higher densities in the dilute phase (Figure 5A). These results show that Hero11-WT-α not only results in a lower density of the concentrated phase of TDP-43 but also increases the population of TDP-43 in the dilute phase. The former suggests essential roles of electrostatic repulsion between Hero11-WT-α chains, whereas the latter implies contributions from attractive interactions between Hero11-WT-α and TDP-43. The coexistence of both repulsive and attractive interactions complicates the net effect of Hero11 on TDP-43’s condensation and requires a more detailed analysis as discussed below.

Since the density changes of TDP-43 induced by Hero11-WT-α are the most significant at 290K (Figure 5A), we will focus on this temperature in the following discussion. To decipher the regulatory function of Hero11 on TDP-43’s condensation at the molecule level, we analyze the number of contacts formed by each chain of TDP-43. We performed five more independent simulations for every 100 TDP-43 + 10*n* Hero11-WT-α (*n* = 1,2, …, 9) system to get more sampling. Each simulation was run for 9× 10^7^ steps, and the last 3 × 10^7^ steps were used for data analysis. At 290K, the dense and dilute phases can be well defined. Thus, we can separately analyze the inter-chain contacts of the proteins in the two phases. Specifically, we count the number of contacts formed by every TDP-43 chain (*c*_TDP − 43_) with the other TDP-43s (TDP-43-TDP-43) or Hero11s (TDP-43-Hero11). Figure 5B shows the average (over all the structures simulated at the same condition) inter-chain contact number of TDP-43 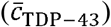 as a function of the number of Hero11-WT-α included in the simulations (*n* _Heor11-WT-α_). As shown in the upper panel of Figure 5B, compared with the 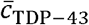 value in the homotypic TDP-43 condensate (the horizontal dotted line, ∼2800), adding Hero11-WT-α to the system diminishes the number of TDP-43-TDP-43 contacts. As *n* _Heor11-WT-α_ increases to 90, the TDP-43-TDP-43 contact number drops to ∼2200 (see the dashed line and empty dots). Apparently, this phenomenon can be explained by the replacement of some TDP-43-TDP-43 contacts by TDP-43-Hero11-WT-α interactions in the condensate. Indeed, as the number of Hero11-WT-α increases, the TDP-43-Hero11 contact number increases, which can be interpreted from the difference between the solid (total) and dashed (TDP-43-TDP-43) lines in the upper panel of Figure 5B. However, our results also show an evident decrease in the total contact number formed by each TDP-43, which cannot be solely explained by substituting TDP-43-TDP-43 contacts with TDP-43-Hero11 contacts. Considering that the repulsive electrostatic interactions result in an extraordinarily low *T*_*C*_ of Hero11-WT-α, it is straightforward to speculate that the repulsion between Hero11-WT-α also induces the decrease of TDP-43’s contact number. To testify to this assumption, we analyze the contact numbers of TDP-43 from the simulations containing 100 TDP-43 and 100 Hero11-KRless-α (results shown in Supplementary Figure S13). We find that when all the positive charges are removed, instead of reducing TDP-43’s total contact number, Hero11-KRless-α increases 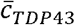 (Supplementary Figure S13), consistent with the higher *T*_*C*_ yielded by Hero11-KRless-α (Figure 4E). These results illustrate the function of Hero11-WT-α in the concentrated phase of TDP-43 that Hero11-WT-α replaces some part of TDP-43’s inter-chain contacts, and the strong electrostatic repulsion between Hero11-WT-α chains “loosen” the whole condensate, resulting in fewer intermolecular interactions on TDP-43.

We also analyzed TDP-43’s contact number in the dilute phase, as shown in Figure 5B’s lower panel. We find that both the total contact number and the number of TDP-43-Hero11 contacts grows as *n*_Hero11-WT-α_ increases. This result suggests that TDP-43 has a higher probability to stay in the dilute phase due to interactions with Hero11-WT-α and explains the increased saturation concentration of TDP-43 in the presence of Hero11-WT-α (see Figure 5A, the low-density branches). We then calculate the radius of gyration of TDP-43 (*R*_*g*,TDP − 43_) in the dilute phase to evaluate the effect of the increased intermolecular contacts on the conformation of TDP-43. As shown in Figure 5C, *R*_*g*,TDP − 43_ expands from ∼31.2Å to ∼31.5Å when *n*_Hero11-WT-α_ increases from 10 to 90. Notably, all these *R*_*g*,TDP − 43_ values are larger than the one calculated in the single-chain simulations (Figure 5C). In contrast, *R*_*g*,TDP − 43_ in the dense phase is independent of the concentration of Hero11-WT-α and close to the value in the homotypic TDP-43 condensation (Supplementary Figure S14). As discussed earlier in this work, TDP-43 in the condensate has larger *R*_*g*_ values than in the dilute phase, indicating an entropy gain during the phase separation (Supplementary Figure S14). Therefore, the results of *R*_*g*,TDP − 43_ in the presence of Hero11-WT-α indicate that more extended structures of TDP-43 in dilute phase are favored, which reduces the conformational entropy differences of TDP-43 between the two phases. Our results show that Hero11-WT-α regulates the thermodynamic behavior of TDP-43 with both energetic (Figure 5B) and entropic (Figure 5C) contributions.

To quantitatively evaluate the effect of Hero11-WT-α on the kinetics of TDP-43, we calculated the diffusion coefficient of TDP-43 (*D*_TDP − 43_) in both dense and dilute phases, respectively. Figure 5D shows *D*_TDP − 43_ in the two phases as functions of *n*_Hero11-WT-α_. Interestingly, the effects of Hero11-WT-α on TDP-43’s diffusion in the two phases are opposite. In the dilute phase, due to the interactions with Hero11-WT-α, TDP-43 is slowed down compared with its diffusion speed in the homotypic condensate (Figure 5D lower panel). In contrast, in the dense phase, TDP-43 diffuses faster when more Hero11-WT-α is involved (Figure 5D upper panel). This result further illustrates that Hero11-WT-α decreases the number of intermolecular interactions of TDP-43 (Figure 5B). Importantly, higher diffusion speed of TDP-43 induced by Hero11-WT-α implies that the condensation formed by TDP-43 and Hero11-WT-α is more fluidic than the homotypic TDP-43 assembly. This result supports the anti-aggregation function of Hero11 [28] with a molecule-level kinetic explanation.

### Effect of Hero11’s secondary structure on its regulatory function

As already shown in this work and previous studies, α-helical secondary structures affect the homotypic phase behaviors of proteins [22,49]. We then ask how heterotypic condensation is affected by the secondary structural properties. To answer this question, we carried out MD simulations for the TDP-43-Hero11-WT-noα mixture. We find that the “noα” systems’ phase diagrams are similar to those with α-helices (see Supplementary Figure S12B). Notably, Hero11-WT-noα also decreases *T*_*C*_ of TDP-43, while Hero11-KRless-noα and Hero11-chargeless-noα do not have this effect (Supplementary Figure S12B). We then calculate the partition coefficient (defined as the number of Heor11 in the dense phase divided by the number of Hero11 in the dilute phase) of Hero11-WT-α and Hero11-WT-noα, respectively, to see the effect of the α-helical secondary structure on the distribution of Hero11 proteins. Interestingly, we find that there are more Hero11-WT-α populated in the dense phase than the Hero11-WT-noα at the same condition (Supplementary Figure S15). This phenomenon is more evident at relatively lower temperatures or at lower Hero11 concentration. Contact map analysis shows that in the mixture of TDP-43 and Hero11-WT-α, a large fraction of inter-molecular contacts between TDP-43 and Hero11-WT-α are contributed by the α-helices from both proteins (Supplementary Figure S16). In the TDP-43-Hero11-WT-noα systems, these secondary structure-facilitated interactions are missing (Supplementary Figure S17). Therefore, the inter-molecular interactions energetically favor Hero11-WT-α in the condensate of TDP-43.

We further analyzed the distribution of Hero11 molecules in the simulation boxes. Figure 6A shows two structures simulated at 260K for 100 TDP-43 in mixture with 90 Hero11-WT-noα or 90 Hero11-WT-α, respectively. These two structures show that there seems to be more Hero11-WT-α than Hero11-WT-noα inside the concentrated phase of TDP-43. To quantitatively assess the distribution of Hero11-WT chains in the simulation boxes, we calculated the time-averaged local density of Hero11-WT-α and Hero11-WT-noα along the *z*-axis (Figure 6B). The density distribution of TDP-43 is also shown, as a reference to indicate the position of the concentrated phase. As can be seen, at both 260K and 290K, the local concentration of Hero11-WT-α chains located in the dense phase is higher than that of Hero11-WT-noα. Additionally, consistent with the structures in Figure 6A, we find that Hero11-WT-noα has high population at the boundary of the TDP-43 condensate, whereas Hero11-WT-α can enter deeper into the dense phase (Figure 6B). These results can be explained by the difference in the secondary structure-dependent inter-molecular contacts (Supplementary Figure S17). As can be seen from the inter-molecular contact map, compared with Hero11-WT-noα, the α-helical regions in Hero11-WT-α contributes more interactions with TDP-43’s α-helical region. These contacts make Hero11-WT-α easier than Hero11-WT-noα to permeate into the interior of TDP-43’s condensation. We find that the difference between Hero11-WT-α and Hero11-WT-noα is weakened at relatively higher temperatures (290K, Figure 6B lower panel). We then calculate the Hero11 concentration-dependent thermodynamic and kinetic quantities of TDP-43 (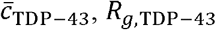, and *D*_TDP − 43_) in the presence of Hero11-WT-noα at 290K and find no substantial difference compared to the results with Hero11-WT-α (see Supplementary Figure S18). Note that in this study, we simulated two extreme structures of Hero11, one for AlphaFold2-predicted one and another for totally disordered one. Based on the atomistic MD simulation results, Hero11 might have weak secondary structure propensity (Figure S2). Therefore, the real structural effect of Hero11 would be between Hero-WT-α and Hero-WT-noα.

**Figure 6.**
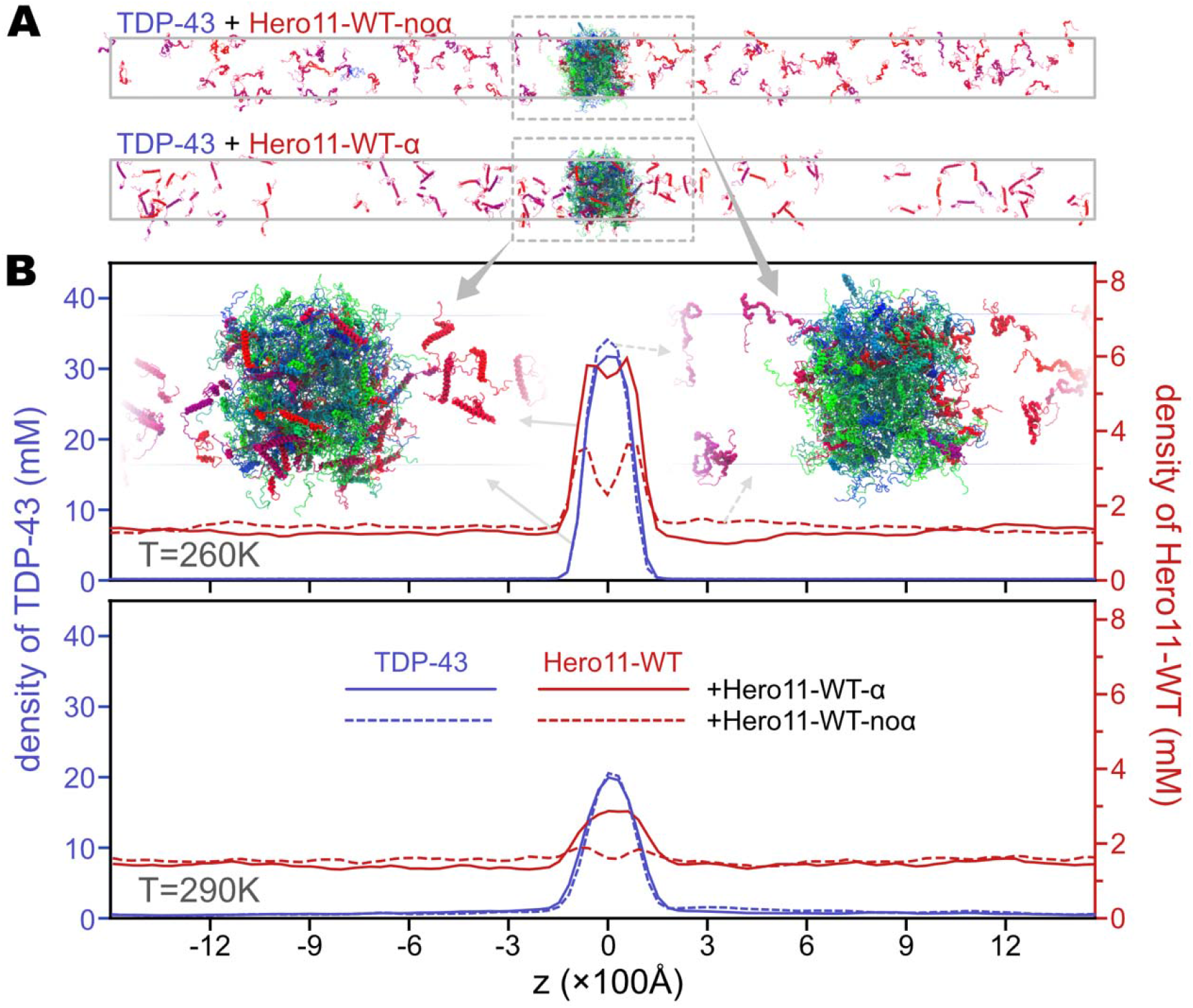
Effect of α-helical secondary structures on the distribution of Hero11-WT in the condensate. (A) The final structures of two simulations at *T* = 260K including 100 TDP-43 + 90 Hero11-WT-noα (upper) or 90 Hero11-WT-α (lower). (B) Densities of TDP-43 (blue) and Hero11-WT (red) along the *z*-axis in simulations at *T* == 260*K* (upper) and 290K (lower), respectively. Solid and dashed lines represent results analyzed from simulations with Hero11-WT-α and Hero11-WT-noα, respectively.

## Discussion

In the current work, we used Hero11 as an example to study the mechanisms of regulation of biomolecular condensation. We find that Hero11 binds to the condensate formed by TDP-43 (Figure 4A-4D), which confirms well to the “scaffold and client” (or “scaffold and ligand”) paradigm [32,60,61]. The stickers-and-spacers model has been used to show that formation or dissolution of the condensate can be regulated by the valency and binding specificity of the client [32]. However, charges and repulsive interactions were not considered in the model [32]. Here, our results demonstrate that Hero11 destabilizes TDP-43’s condensate by introducing strong client-client electrostatic repulsion in addition to the normally attractive client-scaffold interactions. By analyzing the contact number of TDP-43, we show that Hero11 binds with TDP-43 in both dense and dilute phases (Figure 5B, and Supplementary Figures S16, S17, and S18A), but effects of Hero11 are different in the two phases, as explained in the following paragraphs.

Specifically, we consider three possible mechanisms for the Hero11’s regulation of TDP-43’s condensation (Figure 7). First, in the concentrated phase, Hero11 forms contacts with the scaffolding TDP-43 mainly via short-range attractive interactions (modeled by the HPS potential). At the same time, the positively charged residues in Hero11 exerts long-range repulsive forces on Hero11. As a result, Hero11 chains tend to repel each other and, consequently, loosen the whole condensate, including the scaffold they bind on. Accordingly, we observe lower densities (Figure 5A), smaller numbers of contacts (Figure 5B), and higher diffusion coefficients of TDP-43 (Figure 5D) in the dense phase, when Hero11 is present. Second, in the dilute phase, Hero11 increases the total contact number (Figure 5B lower panel, also see Supplementary Figures S16-S18) and the *R*_*g*_ of TDP-43 chains. As discussed earlier, *R*_*g*_ of TDP-43 is larger in the dense phase than in the dilute phase, indicating an entropy gain during the condensation [58,59]. Hero11 results in a larger *R*_*g*_ of TDP-43 in the dilute phase, thus reduces this entropy gain. Therefore, these results explain Hero11’s effects of enhancing the probability of TDP-43 to stay in the dilute phase.

**Figure 7.**
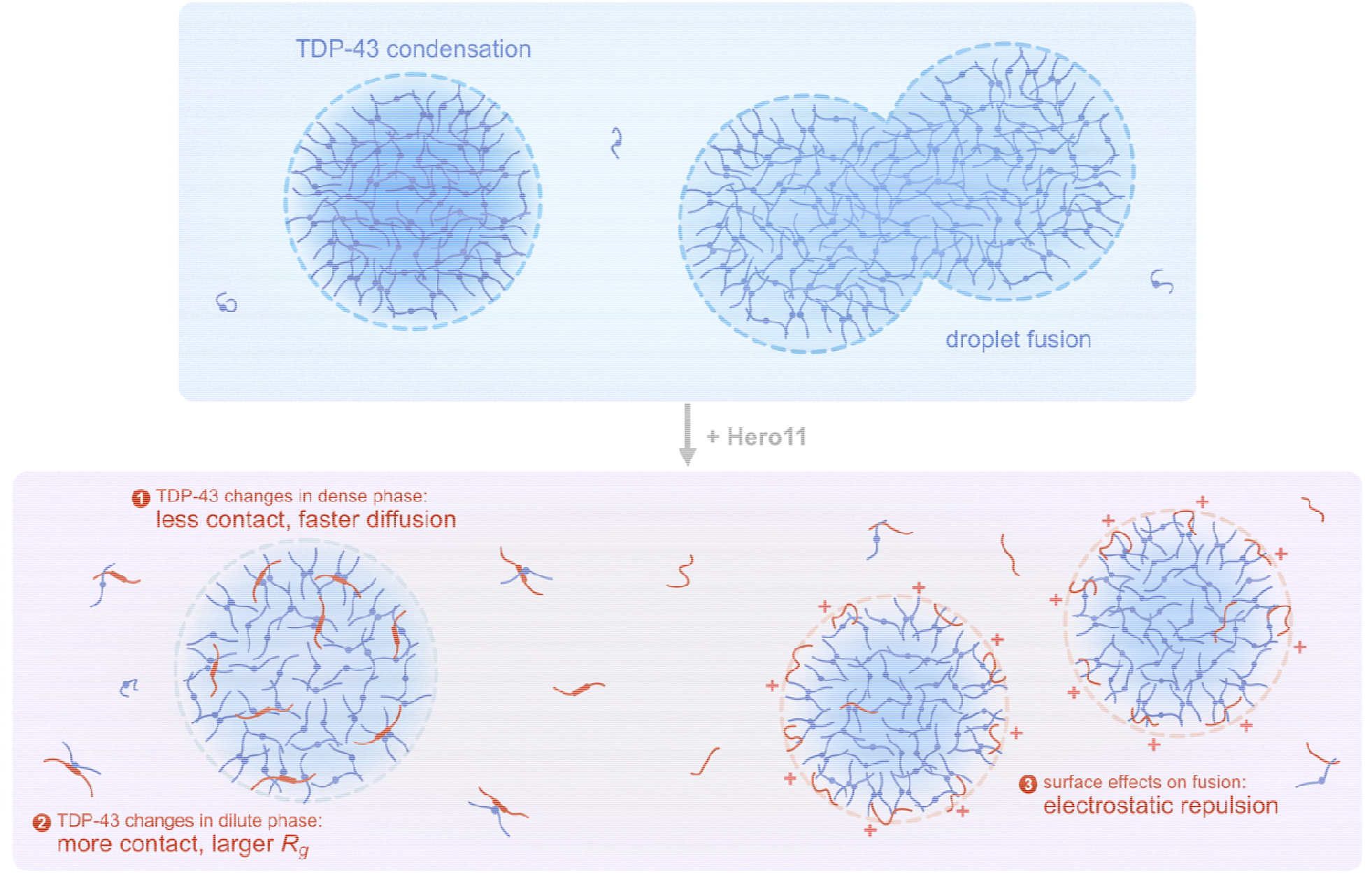
Molecular mechanism of Hero11’s anti-aggregation function. Top: TDP-43 can form homotypic condensation via inter-molecular interactions formed by IDR (lines) and facilitated by α-helical structures (dots) [49]. Bottom: Hero11 regulates TDP-43’s phase behavior using three different mechanisms: 1) Hero11 joins TDP-43’s condensate as a client and decreases the inter-molecular contacts of TDP-43. Consequently, TDP-43’s diffusion is promoted. 2) Hero11 raises the probability for TDP-43 to be in the dilute phase and induces larger of TDP-43 outside of the condensate. 3) In the case of less helical structures, Hero11 tends to stay at the surface of the droplets, which may contribute to reducing fusion [64].

We also show that as the structural helicity decreases, Hero11 distributes more on the surface layer of the co-assembly with TDP-43. Similar results were reported in a previous MD simulation study of the heterotypic condensate formed by LAF-1 and RNA [47]. Notably, amphiphilic proteins have been found to form a surface layer that can regulate the structure and size of condensates [63]. Another combined computational and experimental work reveals that surface charges stabilize and protects biomolecular condensates against fusion [64]. These observations, in combination with our findings of Hero11-WT-noα’s surface distribution (Figure 6), suggest a third possible mechanism of Hero11’s regulatory function that Hero11 may contribute to sustain the condensate of TDP-43 and prevent droplet coalescence (Figure 7). We propose these possible mechanisms with detectable quantities such as radius of gyration and diffusion coefficients to be verified by in vitro and in vivo experiments [65,66].

Our results show that the net charge is dominant in determining the thermodynamic property and regulatory function of Hero11. Similar to Hero11, large absolute value of net charge, either positive or negative, is a common feature of Hero proteins [28]. Therefore, we speculate that the driving force for th other Hero protein’s anti-aggregation function may also be the strong electrostatic repulsion. Interestingly, RNA, which is also highly charged, has also been shown to induce concentration-dependent reentrant phase behaviors of many RNA-binding proteins [27,47,67,68]. Particularly, one recent computational study shows that a high concentration of RNA can slow down the ageing of condensates formed by proteins composed of β-sheet secondary structures [68]. Although the computational models and target systems are different, the results of RNA’s deceleration effect are highly consistent with Hero11’s anti-aggregation mechanism proposed here. In summary, these results suggest that adding repulsive interactions to the framework of the sticker-spacer or scaffold-ligand models can extend their applicability to describe formation, regulation, or destruction of biomolecular condensation.

## Models and Methods

### All-atom simulations of wild-type and KR-less Hero11

We used AlphaFold2 [50] to predict the structure of Hero11 and got two α-helical regions at residues 17-25 and 38-72. To test the stability of these secondary structures, we performed all-atom MD simulations of the wildtype Hero11 and the KR-less mutant with CHARMM36m force field [38]. More detailed information of the atomistic simulations can be found in the Supplementary Methods.

### Coarse-grained modeling of TDP-43 and Hero11

In the CG simulations, we study the full-length Hero11 (99 amino-acid residues) and the C-terminal fraction of TDP-43 (TDP-43, index 261-414). To understand the effect of the charged residues, we also simulate several mutants of Hero11, including the KR-less (all Arginine and Lysine mutated to Glycine), the charge-less (all charged residues mutated to Glycine), and the “scrambles” (sequence randomly permuted). Sequences of all the simulated proteins can be found in Supplementary Table S1.

We employ the HPS CG model for the IDRs [40]. The HPS model represents every amino acid residue as one particle. The potential energy function is given by [40]:

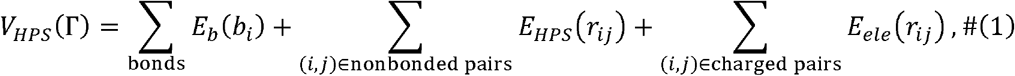

where Γ is the conformation of an IDP, *E*_*b*_ (*b*_*i*_)is a harmonic potential for every two neighboring CG particles with bond length *b*_*i*_:

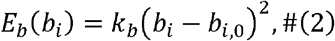

where the reference value of bond length is *b*_*i*,0_ =3.8Å and the force constant is *k*b = 2.39 *kcal* Å^−2^. *E* _*HPS*_ (*r*_*ij*_) is the interaction between two non-bonded particles and is defined by [40,69]:

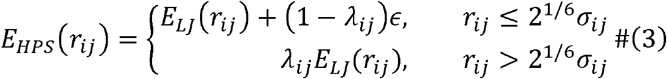

where *λij* is the hydropathy and *E*_*LJ*_(*r*_*ij*_) is the Lennard-Jones potential:

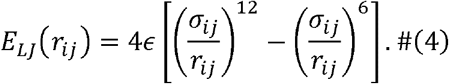

In both equations (3) and (4), *∈* =0.2kcal/mol We use *σ*_*ij*_ values in the original HPS model, and the *λ*_*ij*_ values in the recently proposed HPS-Urry optimization [44]. Both *σ*_*ij*_ and *λ*_*ij*_ use an arithmetic combinational rule for the interacting pair of particles (*i, j*).

As for the electrostatic interaction *E*_*ele*_ (*r*_*ij*_) in Equation (1), we use the Debye-Hu□ckel term:

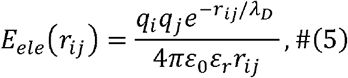

where *r*_*ij*_ is the distance between the nonbonded charged particles, *λ*_*D*_ the Debye screening length, and *ε*_0_ the dielectric permittivity of vacuum. *ε*_*r*_ is the relative permittivity of the solution and is a function of the solution temperature *T* and salt molarity *C*: *εr* = *e*(*T*)*a*(*C*),where *e*(*T*)= 249.4-0.788 *T* + 7.20 × 10^−4^*T*^2^[70], and *a*(*C*) = 1-0.2551*C*+5.151×10^−2^ *C*^2^ -6.889×10^−3^ *C*^3^ [71]. The Debye length is given by 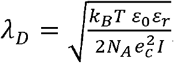 where *N*_*A*_ is the Avogadro’s number, *e*_*c*_ the elementary charge, and *I* the ionic strength of the solution.

As described above, there are α-helical secondary structural pieces in the AlphaFold2 predicted Hero11’s conformation. However, our atomistic simulations did not provide a convergent conclusion of the stability of these secondary structures (Supplementary Figure S2). Therefore, in the CG simulations, we model two extreme cases of Hero11, namely, one with well-folded α-helical model (Hero11-WT-α), and the other as pure IDR (Hero11-WT-noα). We use the AICG2+ model [57] to maintain these folded parts of protein. The AICG2+ potential energy function has the following form [57]:

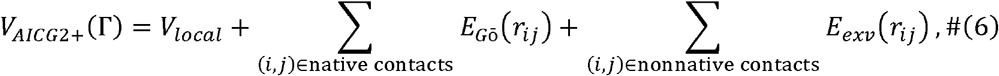

where *V*_*local*_ includes all the bonded terms, *E*_*G*ō_ (*r*_*ij*_) is the Gō-type potential biasing toward the native structure, and *E*_*exv*_ (*r*_*ij*_) is excluded volume interaction. More details of the potential functions can be found in the Supplementary Methods. Specifically, residues 17-25 and 38-72 in Hero11, and residues 60-74 in TDP-43 (320-334 in full-length TDP-43) were modeled with the AICG2+. We use the AlphaFold2-predicted structures as the reference structures for the structure-based potentials. We also turn off the HPS potentials for those non-local particle pairs who already have AICG2+ interactions.

### Coarse-grained molecular dynamics

We carried out all the CG simulations with MD package GENESIS v1.7.1 [72–74]. Structure and topology files were prepared with GENESIS-CG-tool [74]. Time integration step size of the CG simulations is 10fs. Nonlocal interactions *E*_*HPS*_ and *E*_*ele*_ have cutoffs of 20Å and 35Å, respectively.

For both TDP-43 and Hero11, we first simulate single chains for 2 × 10^7^ steps at different temperatures ranging from 110K to 350K. For each protein, we simulate two conformations, one with α-helical region modeled by the AICG2+, the other with pure IDR. The final structures of the single-chain simulations are then duplicated to construct multiple-chain systems, in which we set a minimum distance of 10Å between neighboring chains. We then simulate homotypic systems, including 100 chains of TDP-43 or Hero11, respectively. As for the heterotypic systems consisting of TDP-43 and Hero11, we fix the number of TDP-43 to 100 and add different number of Hero11 into the system. From all the multiple-chain structures, we perform “shrinking” simulations to gradually squash the simulation boxes to 18*nm* × 18*nm z* (*z* is system dependent) in 10^6^ steps. We then extend *z* to 200*nm* for the homotypic Hero11 systems and 300*nm* for all the others. These systems are then equilibrated in the NVT ensemble for 2 × 10^7^ steps at different temperatures. Production runs are conducted with Langevin dynamics with a friction coefficient of 0.01*ps*^−1^.

### Data Analysis

To determine the density of protein particles along the longest dimension (*z*), we divide the *z* axis into *n*_*bin*_ =100 bins. The *z* coordinates of the particles are then used to calculate the “local particle density” in each bin as *ρ*_*partical*_ (*i*) = *N*_*partical*_ (*i*)/ *V*_*bin*_ (*i*), where *N*_*partical*_ (*i*) and *V*_*bin*_ (*i*) are the number of particles and volume of the *i*-th bin. The protein density in the *i*-th bin is then computed as *ρ*_*partical*_ (*i*) =*ρ*_*partical*_ (*i*) /*naa,portein*, where *naa,portein*, is the number of amino-acid residues in the considered protein (*n*_*aa*_,TDP-43 =154 and *n*_*aa*_,Hero11 = 99). The slab, which corresponds to the largest cluster of proteins, is determined in two steps. We first find out the largest *ρ*_*partical*_ (*j*) and its bin index *j*. Then, from this local density peak, we scan in the +z and −z directions and determine the boundaries of the slab (*b*_+*z*_ and *b*-*z*) by finding the first bins satisfying *ρ*_*partical*_ (*b*_±*z*_) < *ρ*_*threshold*_. Here, *ρ*_*threshold*_ = 3.4*mM*, which corresponds to roughly 2 proteins in each bin. Practically, we shift the center of the slab (calculated by *b*−*z* + *b*+*z* /2) to the center of the simulation box. The dense phase is then defined as the region in -,, and the dilute phase is defined as the regions [− *n*_*bin*_/2, *b*_−*z*_ − *b*_*buff*_] and [*b*_+*z*_ + *b*_*buff*,_ *n*_*bin*_ /2], where *b*_*buff*_ =2 is a buffering distance from the boundaries of the dense phase. The densities of the dense (*ρ*_*h*_, the “high” density) and dilute (*ρl*, the “low” density) phases are then defined as the averaged densities in the dense and dilute regions. Note that the current method only works for structures containing single slabs. Therefore, we only calculate *ρ*_*h*_ and *ρl* for relatively lower temperatures, where both dense and dilute phases can be well defined. The critical point (*ρ*_*C*_, *T*_*C*_)is computed by fitting the following equations [75]:

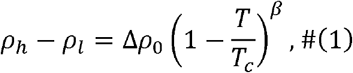

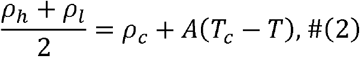

where *ρ*_*C*_ and *T*_*C*_ are critical density and temperature, respectively, Δ*ρ*_0_ and *A* are system-specific fitting parameters, and *β* = 0.325 [76].

To evaluate the interaction between two protein chains, we calculate the number of inter-chain contacts formed by each chain as the following,

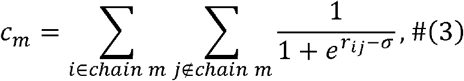

where *C*_*m*_ is the number of contacts of the *m*-th chain, *i* is the index of amino-acid residue in the m-th chain, *j* is the index of residue from all the other chains, *r*_*ij*_ is the distance between the -th and *j*-th residues, and *σ* = 15Å. For TDP-43, we calculate the average number of contacts formed by each chain as 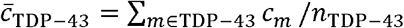, where *n*_TDP-43_ = 100.

To quantitatively describe the diffusion of proteins, we calculated the three-dimensional diffusion coefficient *D*, defined by *D* = MSD(*Δ*t)/(6*Δ*t), where 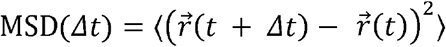 is the mean square displacement of a protein at a time interval of *Δ*t. In Figure 5, the MSD is further averaged over all the proteins in the dense and dilute phases, respectively.

