## Supplementary Information for "Repulsive interaction and secondary structure of highly charged proteins in regulating biomolecular condensation"

### Supplementary Methods

#### Atomistic simulations

We first use AlphaFold2 [1] to predict structures from the sequence of Hero11-WT and Hero11-KRless. Both structures have folded α-helical secondary structures. As a control, we also generate random-coil conformations for these proteins by running coarse-grained simulations and then reconstructing the atomistic structures. These proteins are solvated in TIP3P water boxes. $Na^+$ and $Cl^-$ ions are added to the solution to neutralize the system and raise ionic concentration to 150mM. All the systems have box sizes of ~$181 \times 181 \times 181$Å$^3$ and include 566,884 (Hero11-WT starting from folded), 567,249 (Hero11-KRless from folded), 566,857 (Hero11-WT from random-coil), and 567,198 (Hero11-KRless from random-coil) atoms, respectively.

For each system, we perform energy minimization for 50000 steps and then heat up from 0K to 300K in 100ps. The systems are then equilibrated at 300K in three steps: first 200ps in NPT ensemble with all protein heavy atoms restrained; then 200ps in NPT ensemble with no restraints; and lastly 1050ps in NVT ensemble with no restraints. As production runs, we perform NVT simulations at 300K using Bussi's stochastic velocity rescaling thermostat [2]. CHARMM36m force field [3] is used for energy and force calculations. We set a cutoff of 12Å for van der Waals interactions and use a force-based switch function in the range between 10Å and 12Å. We employ the Particle mesh Ewald (PME) method [4] for electrostatic interactions and set a cutoff of 12Å for the real-space part. The reciprocal-space part is evaluated every two steps. For time integration, we adopt a multiple time scale scheme, r-RESPA [5], with a step size of 3.5fs. All the atomistic simulations are run with GENESIS v2.0 [6–8]. For each system, we carry out 1μs simulations. MD trajectory is stored every 3000 steps (10.5ps).

Secondary structural features of Hero11 are assigned by the Dictionary of protein secondary structure (DSSP) algorithm [9] implemented in the mdtraj package [10].

#### AICG2+ model for protein

As described in the Methods section of the main text, we use AICG2+ model [11] to maintain the structure of the folded parts in TDP-43 and Hero11. The local interaction terms ($V_{local}$ in Eq.6) include the following parts:

$$V_{local} = \sum_{b_i \in \text{bonds}} k_{b,i}\left(b_i - b_{i,0}\right)^2 + \sum_{r_i \in 1-3 \text{ pairs}} -\varepsilon_i \exp\left(\frac{-\left(r_i - r_{i,0}\right)^2}{2w_i^2}\right) + \sum_{\theta_i \in \text{angles}} -k_B T \ln \frac{P_\theta(\theta_i|i)}{\sin\theta_i}$$
$$+ \sum_{\varphi_i \in \text{dihedrals}} -\epsilon_{\varphi,i} \exp\left(\frac{-\left(\varphi_i - \varphi_{i,0}\right)^2}{2\sigma_{\varphi,i}^2}\right) + \sum_{\varphi_i \in \text{dihedrals}} -k_B T \ln P_d(\varphi_i|i)\,, \qquad (S1)$$

where the first term is for bond-length, the second is for every other particles, and the forth is for dihedral angles. The third and fifth terms in Eq.S1 are statistical flexible potentials, where $P_\theta(\theta_i|i)$ and $P_d(\varphi_i|i)$ are residue-type dependent probability distributions of angles and dihedral angles, respectively [12]. $k_B$ is the Boltzmann constant and $T$ is temperature.

$E_{G\bar{o}}(r_{ij})$ in Eq. 6 is given by:

$$E_{G\bar{o}}(r_{ij}) = \varepsilon_{G\bar{o},i,j}\left[5\left(\frac{\sigma_{ij}}{r_{ij}}\right)^{12} - 6\left(\frac{\sigma_{ij}}{r_{ij}}\right)^{10}\right], \qquad (S2)$$

where $r_{ij}$ is the distance between the $i$-th and $j$-th residues, $\sigma_{ij}$ is the value of $r_{ij}$ in the reference structure, and $\varepsilon_{G\bar{o},i,j}$ is the context dependent energy coefficient.

$E_{exv}(r_{ij})$ in Eq. 6 is given by:

$$E_{exv}(r_{ij}) = \begin{cases} -\varepsilon_{exv}\left(\frac{\sigma_{ij}}{r_{ij}}\right)^{12} + \epsilon'_{exv,i,j}, & r_{ij} < r_C \\ 0, & r_{ij} \geq r_C \end{cases} \qquad (S3)$$

where $r_{ij}$ is the distance between the $i$-th and $j$-th residues, $\sigma_{ij}$ is residue-type dependent excluded volume distance [13], $\varepsilon_{exv} = 0.6kcal/mol$, $r_C = 2\sigma_{ij}$ is the cutoff distance, and $\epsilon'_{exv,i,j}$ is used to shift $E_{exv}(r_{ij})$ to 0 at $r_{ij} = r_C$.

To evaluate the stability of the α-helical regions, we describe the "native-ness" (denoted as the $Q$ value) of a structure as the fraction of formed native contacts. A native contact is defined as two residues having any heavy atoms with in 4.5Å from each other in the reference structure. In a given structure, a native contact is considered as "formed" if their $C_\alpha$ distance is within 1.2 times of their native value.

#### Supplementary Figures

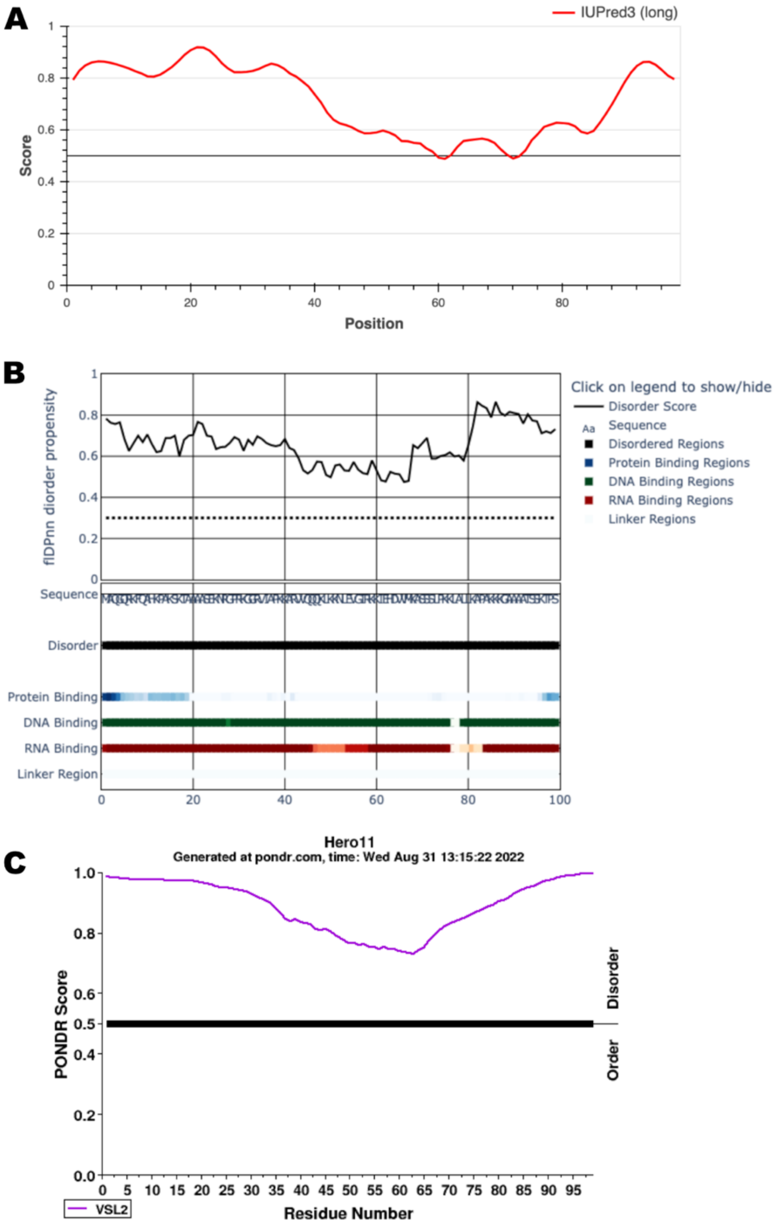


**Figure S1. IDR propensity of Hero11 sequence predicted by IUPred3** [14] **(A), and flDPnn** [15] **(B), and PONDR** [16] **(C) web servers.** Sequence of the wild-type Hero11 can be found from Table S1.

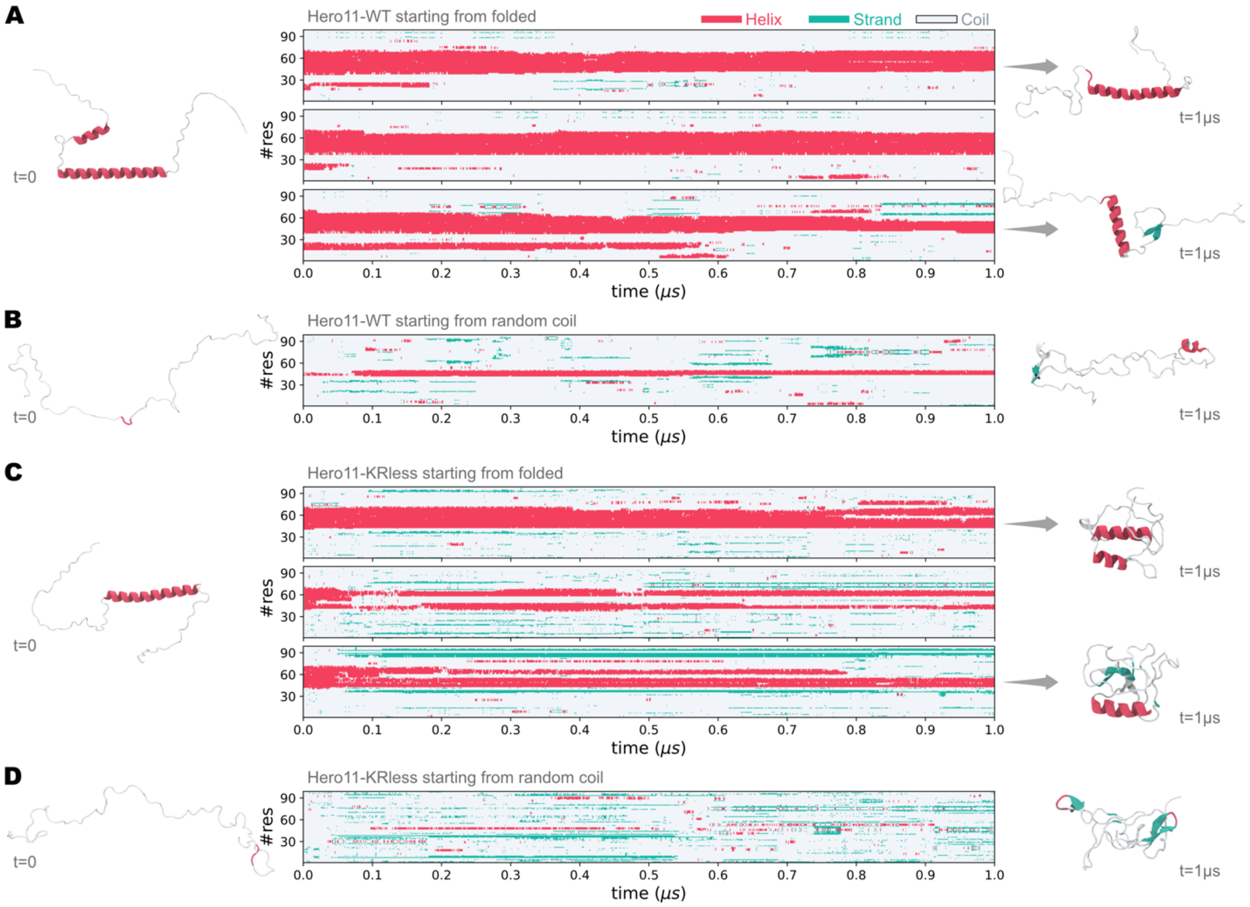


**Figure S2. All-atom simulations of the Hero11-WT and the Hero11-KRless mutant.** (A) Time evolution of secondary structural features of Hero11-WT residues in the simulations started from the AlphaFold predicted structure. Three independent runs were performed and the results are shown in three panels respectively. Secondary structural features are represented by colors, with red for helix (including α-helix, 3/10 helix, and pi-helix), green for β-strand, and gray for coil. The initial structure and two of the final structures are shown on the left and right, respectively. The cartoon structures use the same color code as the secondary structure profile. (B) is the same as (A) but for one simulation of Hero11-WT started from a random coil structure. (C) is the same as (A) but based on three MD trajectories of Hero11-KRless, started from AlphaFold predicted structure. (D) is the same as (B) but for Hero11-KRless initiated from a random coil structure.

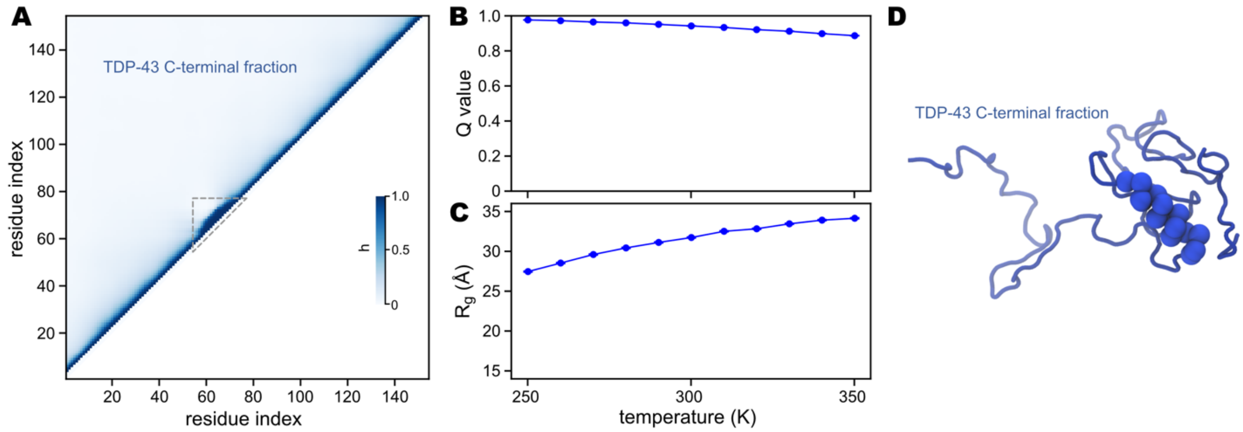


**Figure S3. MD simulations of single-chain TDP-43 C-terminal fraction.** (A) Contact map of TDP-43 analyzed from simulations at 300K. Here, a contact is defined as a function of the distance between two residues: $h_{ij} = 1/(1 + \exp(r_{ij} - 15\text{Å}))$, where $r_{ij}$ is the distance between the $i$-th and the $j$-th residues, and $j > i + 2$. For every non-local pair of residues, we calculated the time average of $h_{ij}$ from 5 independent $2 \times 10^7$-step runs. Value of $h$ is represented by the intensity of blue, as indicated by the color bar. The dashed triangle indicates the α-helical region (indices 60-74). Note that here the residue indices are counted from 1 to 154, which correspond to real indices from 261 to 414 in the full-length TDP-43. (B) Q value (fraction of formed native contacts, upper) and radius-of-gyration ($R_g$, lower) as functions of simulation temperature. Error bars represent the standard errors. (C) A representative structure of TDP-43 simulated at 300K. The α-helical region (indices 60-74) is represented by spheres and the other regions are shown as a thin tube.

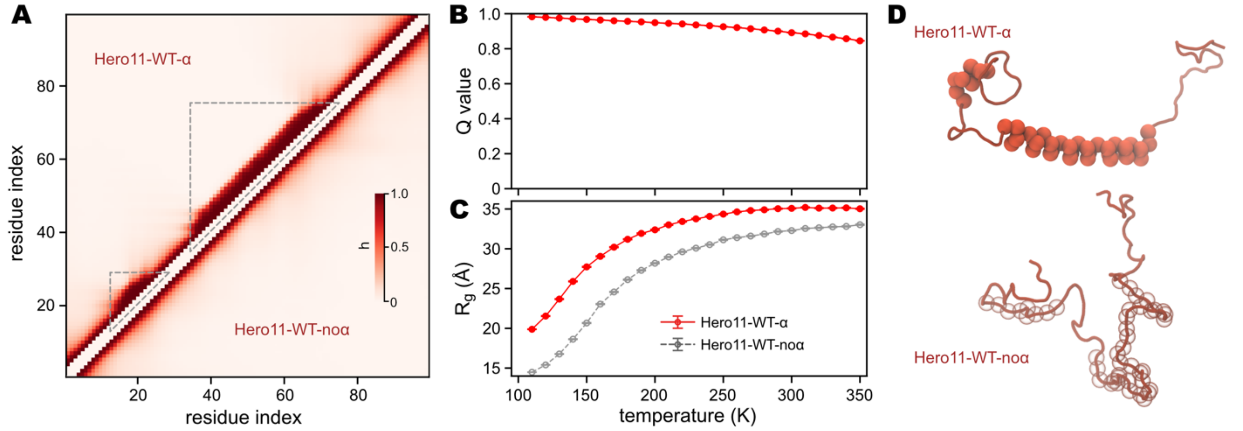


**Figure S4. MD simulations of single-chain Hero11-WT, with or without AICG2+ potentials for the α-helical regions.** (A) Contact map of Hero11-WT-α (upper-left) and Hero1-WT-noα (lower-right), analyzed from simulations at 300K. Same as in Figure S1, a contact is defined as a function of the distance between two residues: $h_{ij} = 1/(1 + \exp(r_{ij} - 15\text{Å}))$, where $r_{ij}$ is the distance and $j > i + 2$. For every non-local pair of residues, we calculated the time average of $h_{ij}$ from 5 independent $2 \times 10^7$-step runs. Value of $h$ is represented by the intensity of red, as indicated by the color bar. The dashed triangles indicate the α-helical regions in Hero11-WT-α (indices 17-25 and 38-72). (B) Q value (fraction of formed native contacts, upper) and radius-of-gyration ($R_g$, lower) as functions of simulation temperature. Error bars represent the standard errors. (C) Representative structures of Hero11-WT with (upper) or without (lower) α-helical region simulated at 300K. The α-helical regions (indices 17-25 and 38-72) are represented by solid spheres. In the model without α-helix, we show the same regions (indices 17-25 and 38-72) as transparent spheres for comparison. All the other regions are shown as thin tubes.

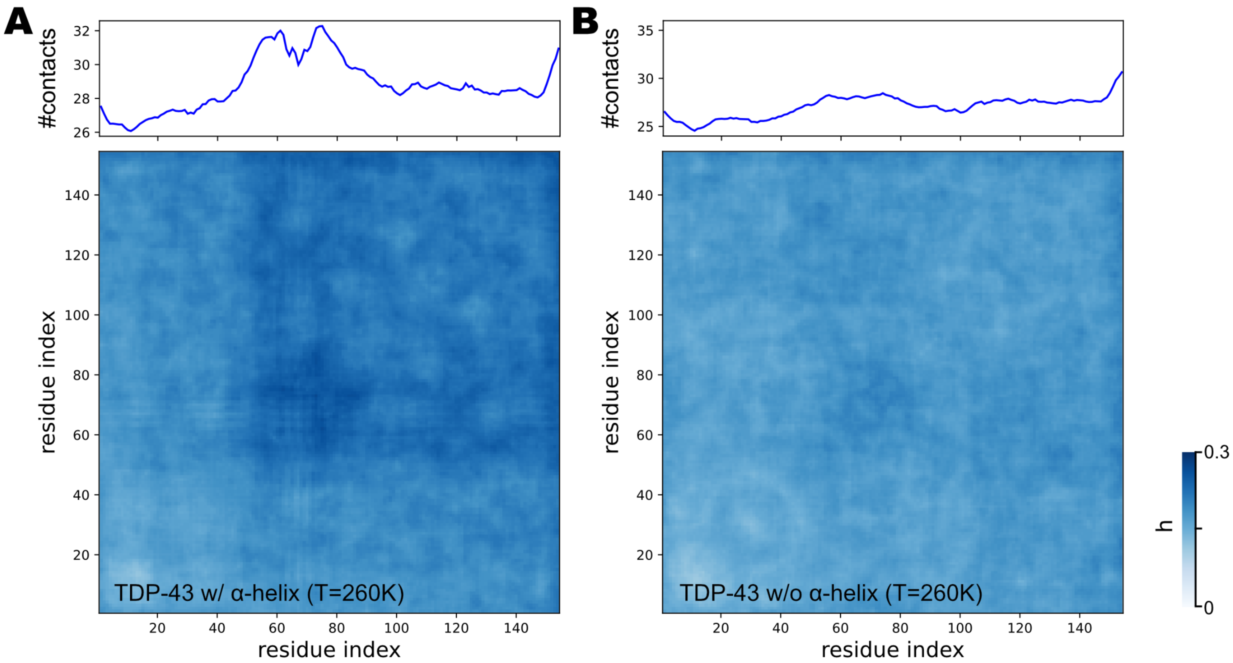


**Figure S5. Inter-chain contact map (bottom) and number of contacts (top) for TDP-43 residues**. These results are calculated from simulations with (A) or without (B) α-helical structure, at temperature 260K. For a pair of residues $i$ and $j$ in a given structure (one frame of the MD trajectory), we count the number of contacts over all the chain pairs: $h_{ij} = \sum_{m<n} \lambda_{ij}(m,n)$, where $\lambda_{ij}(m,n)$ is the contact between residue $i$ from chain $m$ and residue $j$ from chain $n$: $\lambda_{ij}(m,n) = 1/(1 + \exp(r_{ij}(m,n) - 15Å))$. We then calculate the time average of $h_{ij}$ from 5 independent $5 \times 10^7$-step runs (only the latter half of each simulation is used). Value of $h$ is represented by the intensity of blue, as indicated by the color bar. The total number of contacts formed by residue $i$ (top) is then computed as $\sum_j h_{ij}$.

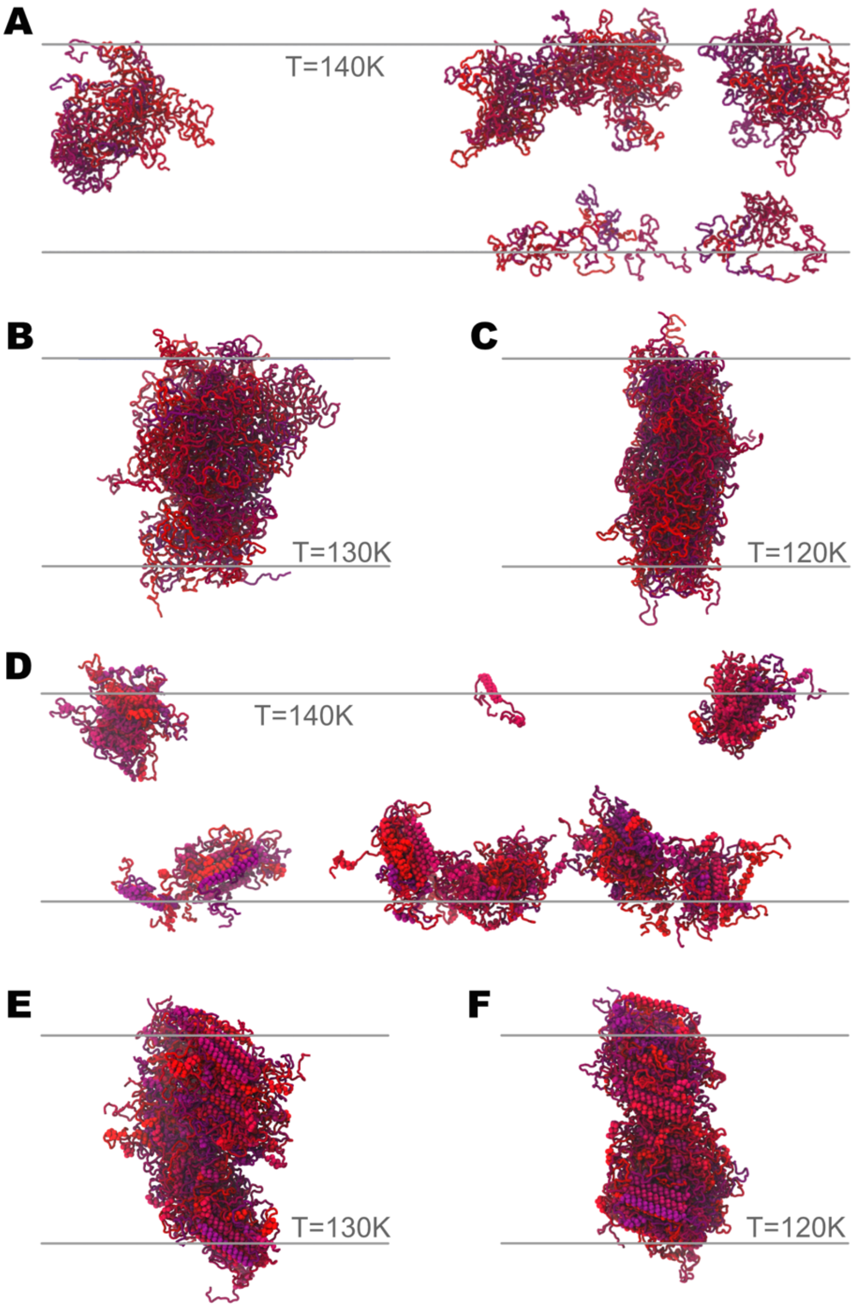


**Figure S6. Representative structures of Hero11-WT-α and Hero11-WT-noα.** (A-C) Structures of Hero11-WT-noα simulated at 140K (A), 130K (B), and 120K (C), respectively. For clarity, only part of each simulation box is shown: $\Delta z = 300$Å for (A) and (B), and $\Delta z = 700$Å for (C). (D-F) are the same as (A-C) but for Hero11-WT-α.

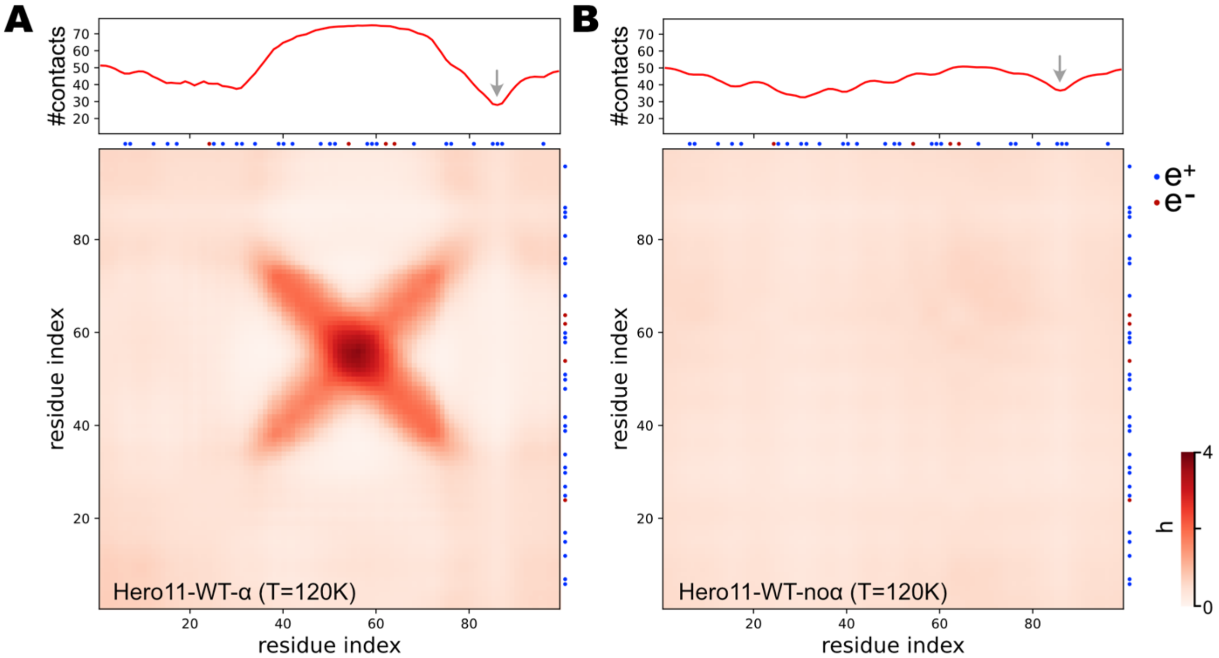


**Figure S7. Inter-chain contact map and total number of contacts for Hero11-WT residues**. These results are based on simulations of Hero11-WT-α (A) and Hero11-WT-noα (B) at temperature 120K. Contact has the same definition as Figure S5. Small dots next to the top and right axes of the contact map indicate the position of the positively (blue) and negatively (red) charged residues, respectively. The gray arrows in the top panels indicate the position of the $K_{85}K_{86}K_{87}$ sequence piece.

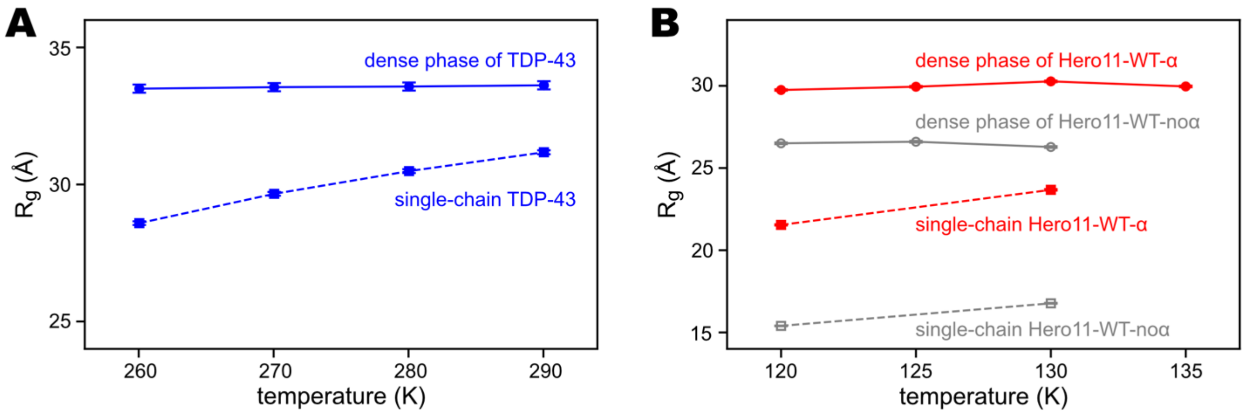


**Figure S8. Radius of gyration ($R_g$) of TDP-43 (A) and Hero11-WT (B) in multi-chain and single-chain simulations.** The single-chain simulation results are the same as shown in Figure S3 and S4. As for the multi-chain simulations, we calculate the average $R_g$ in the dense phase at temperatures below the $T_C$s.

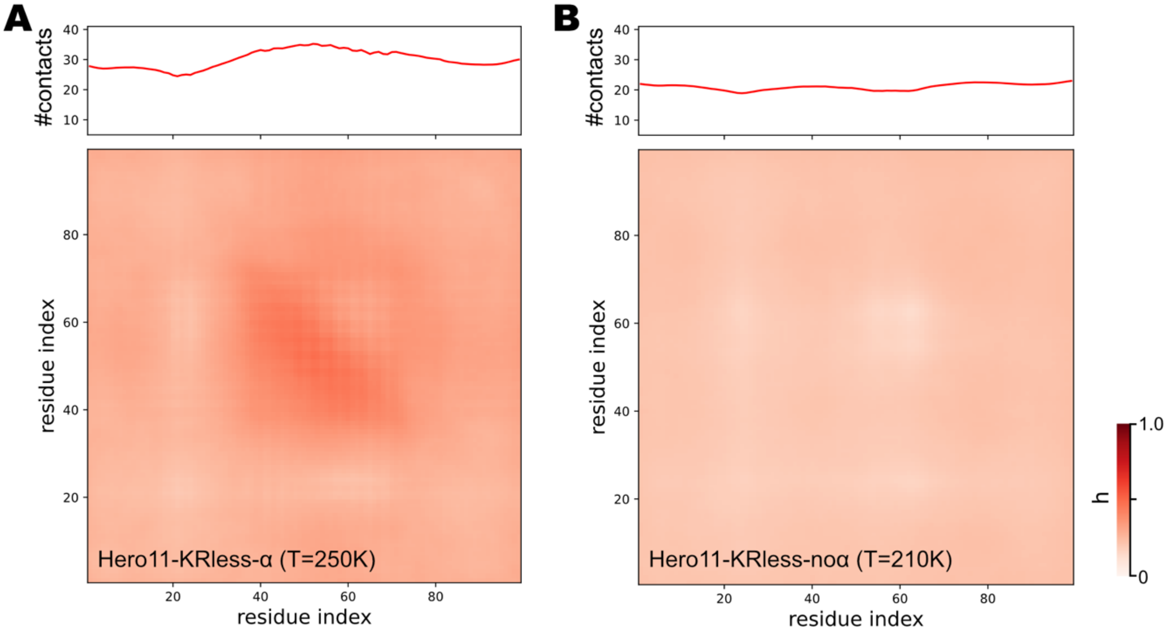


**Fig S9. Inter-chain contact map and number of contacts for residues of Hero11-KRless**. These results are based on simulations of Hero11-KRless-α (A) and Hero11-KRless-noα (B). Contact definitions are the same as Figure S5.

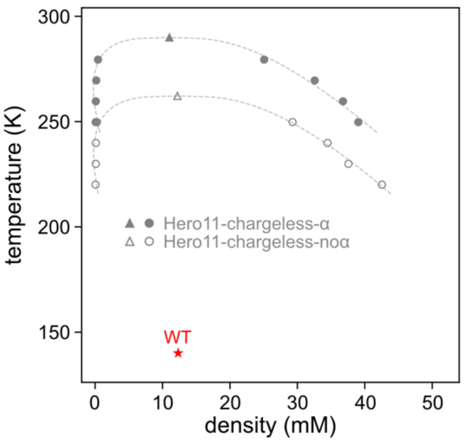


**Figure S10. Phase diagram of the Hero11-chargeless mutant**. In the chargeless mutant, all the charged residues (Arg, Lys, Asp, and Glu) are mutated to Gly. The red star represents Hero11-WT-α's critical point.

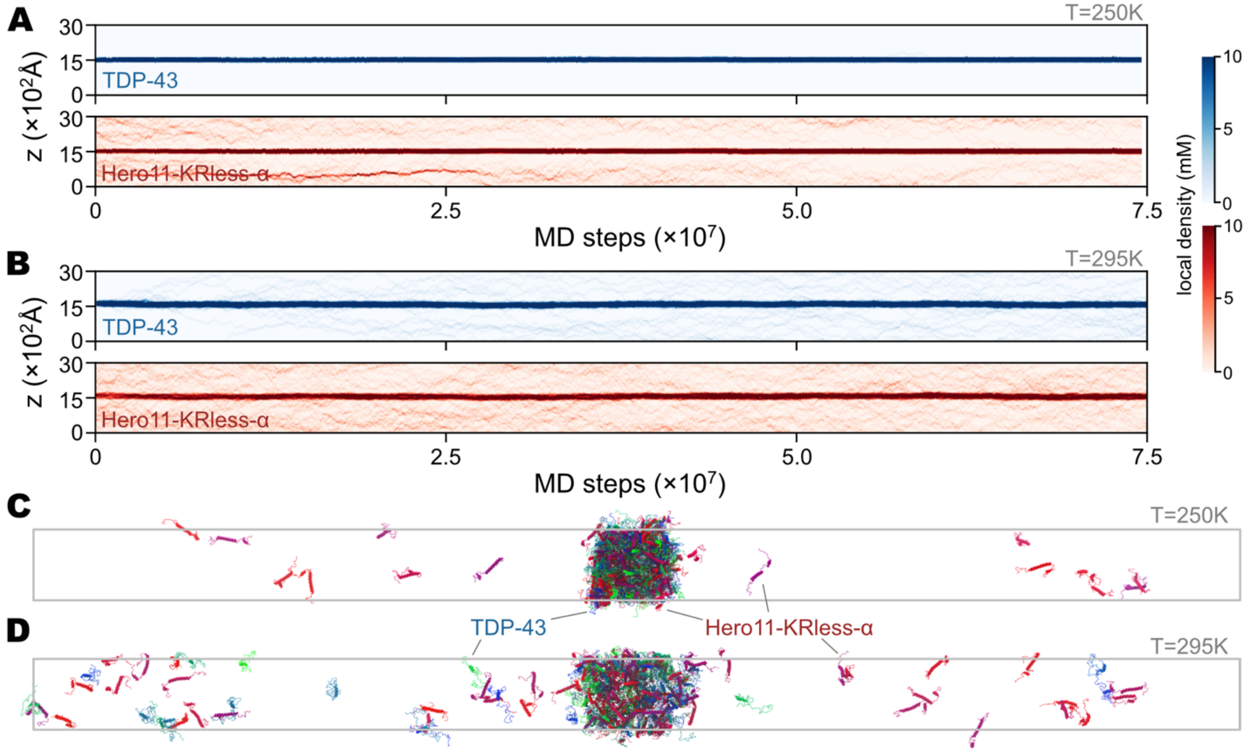


**Figure S11. Slab simulations of heterotypic systems consisting of 100 TDP-43 and 100 Hero11-KRless-α.** (A) and (B) show graphs of local densities of TDP-43 and Hero11-KRless-α simulated at 250K and 295K, respectively. Intensity of blue (red) represents local density of TDP-43 (Hero11-KRless-α), as indicated by the color bars. (C) and (D) show the last structures of the 250K and 300K simulations, respectively. TDP-43 chains are shown in green-blue colors, while Hero11-KRless-α chains are shown in red-purple colors.

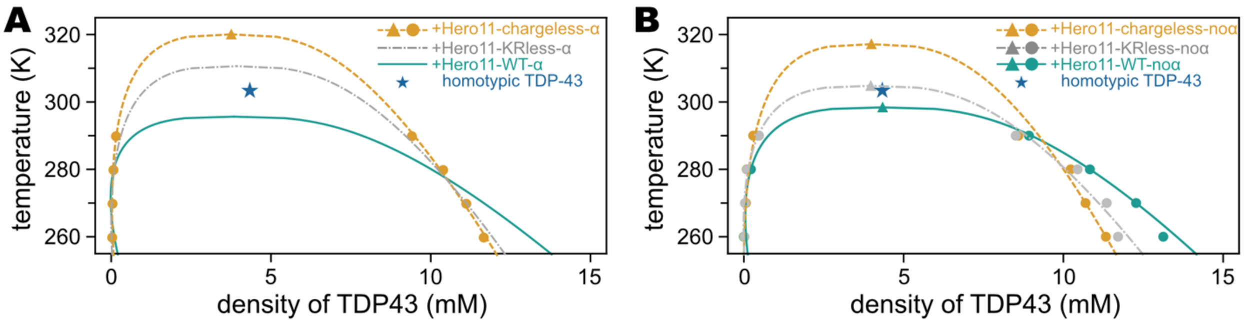


**Figure S12. Full phase diagram of TDP-43 in the presence of wild-type and mutated Hero11s.** (A) Phase diagram of TDP-43 with or without Hero11 (WT and mutants). Hero11-WT-α and Hero11-KRless-α data is shown in cyan (solid) and gray (dot-dashed), which are the same as shown in Figure 4 in the main text. Circles represent the densities calculated from the simulations, whereas triangles are the fitted values (see Methods). The blue star indicates the critical point of homotypic TDP-43. (B) is the same as (A) but for simulations of TDP-43 and Hero11 models without α-helical secondary structures.

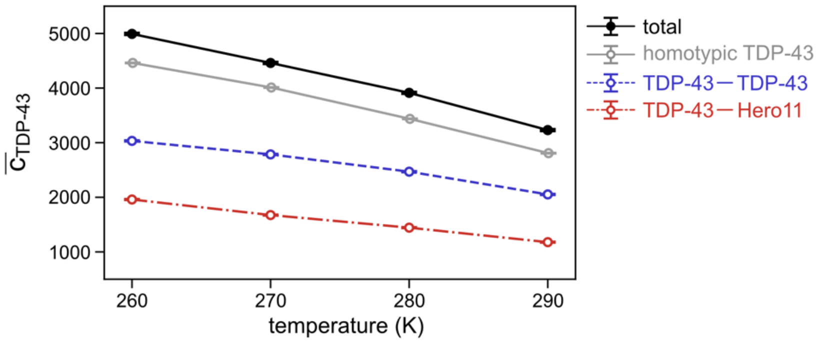


**Figure S13. Average number of inter-chain contacts formed by TDP-43 chain ($\bar{c}_{\mathbf{TBP-43}}$) in the simulations containing 100 TDP-43 and 100 Hero11-KRless-α.** The total number of contacts (black) is the sum of TDP-43-TDP-43 (blue dashed line) and TDP-43-Hero11 (red dashed line) contacts. Gray line is $\bar{c}_{\mathrm{TBP-43}}$ calculated from the homotypic TDP-43 simulations.

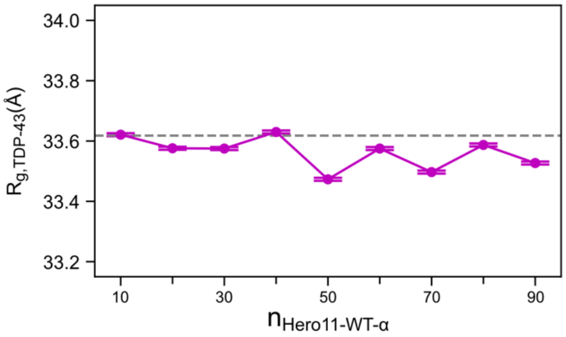


**Figure S14. Radius of gyration of TDP-43 ($R_{g,\mathbf{TDP-43}}$) in the dense phase, as a function of the number of Hero11-WT-α included in the simulation ($n_{\mathbf{Hero11-WT-\alpha}}$).** Note that the difference between the maximum and minimum of $R_{g,\mathrm{TDP-43}}$ is less than 0.2Å, much smaller than the one observed in the dilute phase (Figure 5C).

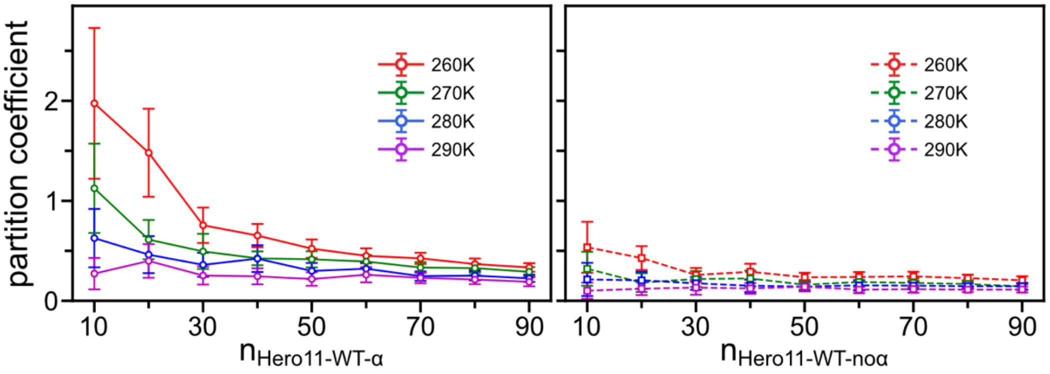


**Figure S15. Partition of Hero11-WT-α and Hero11-WT-noα in and out of the TDP-43 condensation.** Partition coefficient (fraction of Hero11 in the dense phase divided by fraction of Hero11 in the dilute phase) as a function of total number of Hero11 in the simulation system. Left: TDP-43 + Hero11-WT-α; Right: TDP-43 + Hero11-WT-noα.

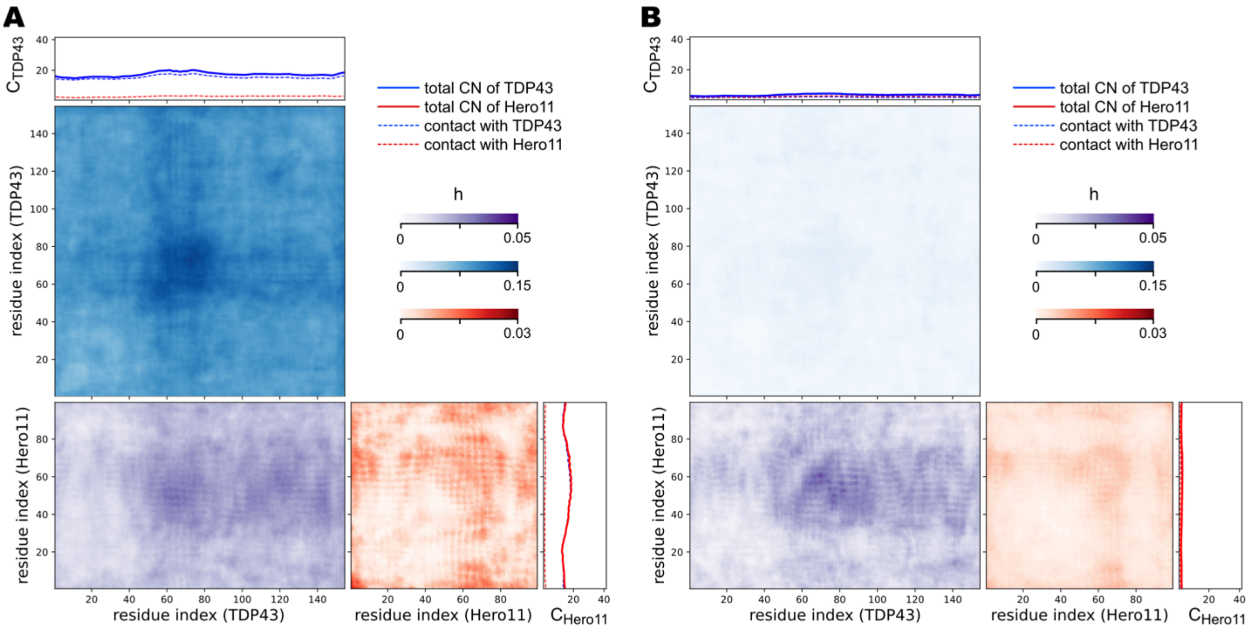


**Figure S16. Inter-molecular contact map for TDP-43 and Hero11-WT-α residues in simulations of 100 TDP-43 + 90 Hero11-WT-α at 290K.** (A) and (B) are for the dense and dilute phases, respectively. TDP-43-TDP-43, Hero11-Hero11, and TDP-43-Hero11 contacts are colored in blue, red, and purple, respectively. Intensity of color indicates the contact number using the same definition as Figure S5. Here, TDP-43-TDP-43 contacts (blue) are normalized by the number of TDP-43, Hero11-Hero11 contacts (red) are normalized by the number of Hero11, whereas TDP-43-Hero11 contacts (purple) are normalized using the number of TDP-43.

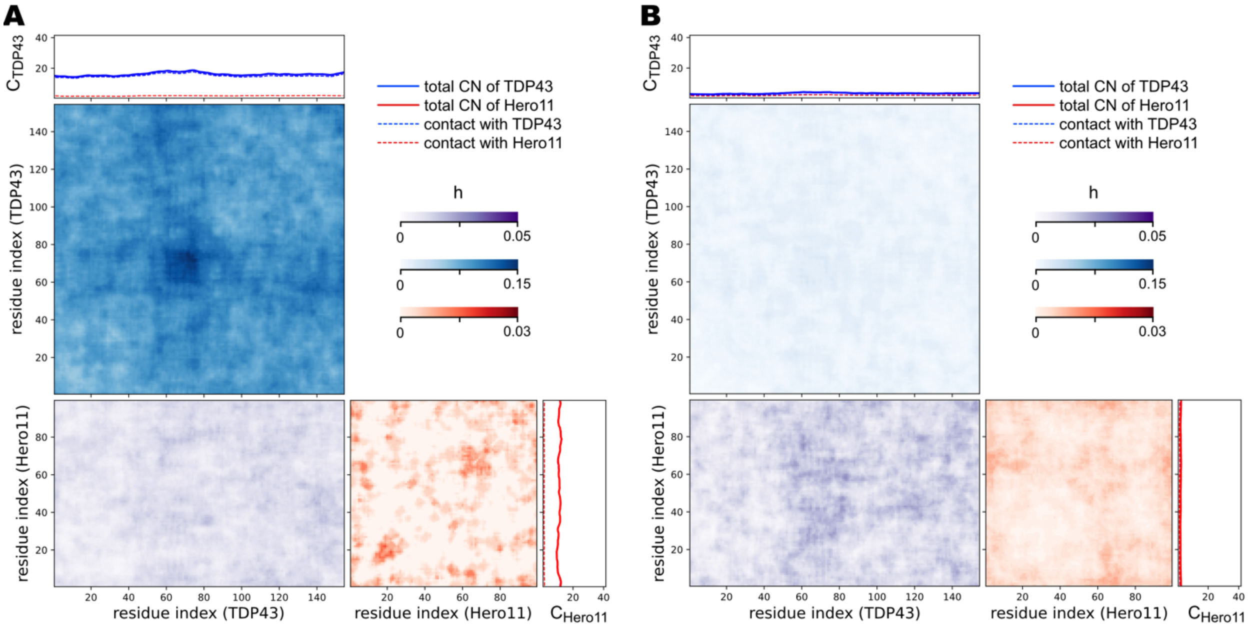


**Figure S17. Inter-molecular contact map for TDP-43 and Hero11-WT-noα residues in simulations of 100 TDP-43 + 90 Hero11-WT-noα at 290K.** (A) and (B) are for dense and dilute phases, respectively. Color scheme is the same as Figure S16.

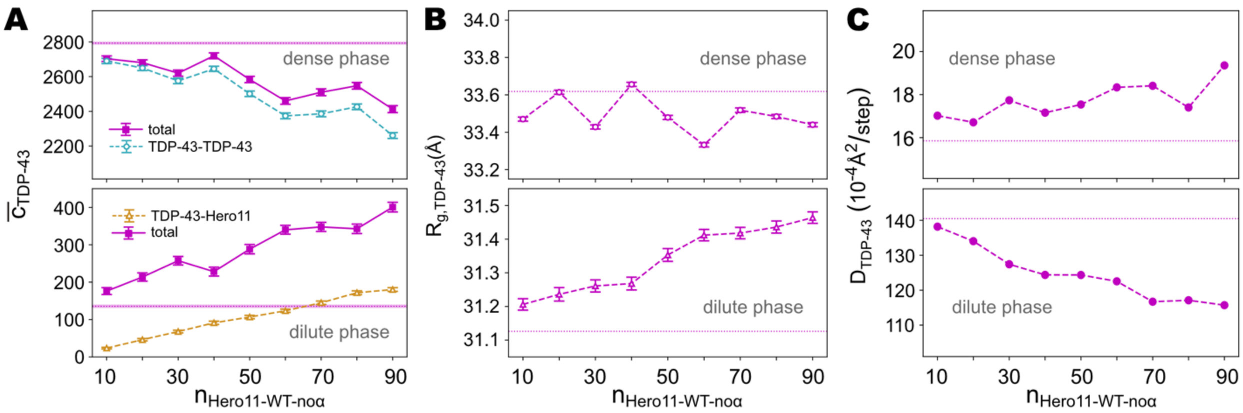


**Figure S18. Contact number, radius of gyration, and diffusion coefficient of TDP-43 as functions of number of Hero11-WT-noα included in the simulations.** (A) Contact number formed by TDP-43 ($\bar{c}_{\mathrm{TDP-43}}$) in the dense (upper) and dilute (lower) phases as functions of the number of Hero11-WT-noα. The total number of $\bar{c}_{\mathrm{TBP-43}}$ (purple) is the sum of TDP-43-TDP-43 (cyan) and TDP-43-Hero11 (orange) contacts. For clarity, in the dense phase we don't show TDP-43-Hero11 data, while in the dilute phase TDP-43-TDP-43 is hidden. (B) Radius of gyration of TDP-43 ($R_{g,\mathrm{TDP-43}}$) in the dense (upper) and dilute (lower) phases as functions of the number of Hero11-WT-noα. (C) Diffusion coefficient of TDP-43 ($D_{\mathrm{TDP-43}}$) in the dense (upper) and dilute (lower) phases as functions of the number of Hero11-WT-noα. All the horizontal dotted lines show the corresponding value calculated from the homotypic TDP-43 system.

**Supplementary Table S1: Sequences of simulated proteins.**

| Protein name | Sequence |
|---|---|
| Hero11-WT | MAQGQRKFQA HKPAKSKTAA AASEKNRGPR KGGRVIAPKK ARVVQQQKLK |
| | KNLEVGIRKK IEHDVVMKAS SSLPKKLALL KAPAKKKGAA AATSSKTPS |
| Hero11-KRless | MAQGQGGFQA HGPAGSGTAA AASEGNGGPG GGGGVIAPGG AGVVQQQGLG |
| | GNLEVGIGGG IEHDVVMGAS SSLPGGLALL GAPAGGGGAA AATSSGTPS |
| Hero11-chargeless | MAQGQGGFQA HGPAGSGTAA AASGGNGGPG GGGGVIAPGG AGVVQQQGLG |
| | GNLGVGIGGG IGHGVVMGAS SSLPGGLALL GAPAGGGGAA AATSSGTPS |
| Hero11-scramble-1 | KPKQAKLSEK SSFQAPAVGP LVKRRENARR AKAVTKLQEK KHIAKTVMKS |
| | VKKKLIASKN TASMSKRAQA SPKQGLVKGQ HARAPKAIKA ALKGGPAKD |
| Hero11-scramble-2 | APSKVKQRIV PKNKSAAKKP FSSDAVKRKS QKERAEMTGV KEGAKLKKAL |
| | AAKPLIKPAP MKKATAAKNR AHIAVKVKAK HGSGQGQTLQ KLRLGSRAQ |
| Hero11-scramble-3 | MSPKQKRTSA KVKAVGILAK LKPSASQLAH NSAKKAEKRV PLIKVTIGPQ |
| | LAGKAQSDVN MLRSAKGAKK VKEKPFKARG AKSKQTAHPR KAKEGRKAA |
| TDP-43 (C-terminal domain) | EPKHNSNRQL ERSGRFGGNP GGFGNQGGFG NSRGGGAGLG NNQGSNMGGG |
| | MNFGAFSINP AMMAAAQAAL QSSWGMMGML ASQQNQSGPS GNNQNQGNMQ |
| | REPNQAFGSG NNSYSGSNSG AAIGWGSASN AGSGSGFNGG FGSSMDSKSS |
| | GWGM |
